# Circadian misalignment underlies immune escape in breast cancer

**DOI:** 10.64898/2026.05.26.726543

**Authors:** Chuqian Liang, Zunpeng Liu, Xiaoke Xu, Zhen Xu, Qi Zhang, Tao Zhang, Riley J. Mangan, Bingqiu Xiu, Gongwei Wu, Tara Akhshi, Zachary M. Sandusky, G. Kenneth Gray, Na Zhang, Genevra Kuziel, Nicole A. Traphagen, Stuart Benjamin Fass, Amy Grayson, Li-Lun Ho, Shengqing Stan Gu, Song Wang, Zion Zihan Sheng, Yi Zhang, Elio Adib, Tianlong Chen, Rinath Jeselsohn, Manolis Kellis, Myles Brown

## Abstract

Circadian regulation shapes tissue physiology, yet how it organizes cellular and molecular dynamics within the tumor microenvironment (TME) and influences tumor immunity remains unclear. Using temporal single-nucleus multiomic profiling of mouse breast tumors, we uncovered extensive circadian programs that are both cell-type-specific and shared across the TME, governing proliferation and immune responses. Notably, cancer epithelial cells exhibited global acrophase misalignment relative to immune populations. This intercellular desynchrony manifests as temporal decoupling between tumor proliferation and immune activation, discordant antigen presentation and T cell recognition with intrinsic activation–exhaustion overlap in T cells, and asynchronous PD-1/PD-L1 oscillations that sustain checkpoint-mediated suppression. Similar patterns were observed in human triple-negative breast cancer (TNBC). Together, these findings establish intercellular circadian misalignment as a mechanism of tumor immune evasion and position the circadian architecture of the tumor–immune ecosystem as a previously underappreciated determinant of tumor development and therapeutic response.

**HIGHLIGHTS:**

- Single-cell multiomics maps circadian regulation of tumor–immune programs.
- Circadian regulation in cancer epithelial cells is misaligned with immune cell populations.
- Tumor–immune temporal misalignment undermines antitumor immunity.
- Circadian misalignment in human TNBC suggests relevance for immunotherapy timing.

## INTRODUCTION

Circadian rhythms are endogenous 24-hour oscillations that temporally organize physiological processes across tissues and cellular systems.^1–5^ Within tumors, circadian programs influence cancer cell proliferation and survival, while also regulating immune cell trafficking, cytokine production, and antitumor effector functions in the tumor microenvironment (TME).^6–8^ Disorganized circadian regulation is associated with enhanced tumor initiation, accelerated progression, and increased metastatic potential.^9–12^ Recent clinical studies, including randomized trials in non–small cell lung cancer and melanoma, have shown that the efficacy of cancer immunotherapy may depend on the time of administration, underscoring the therapeutic relevance of circadian regulation in oncology.^13–15^ Collectively, these findings position circadian regulation as a fundamental dimension of tumor biology and therapeutic responsiveness, underscoring the need to incorporate temporal dynamics into our understanding and treatment of cancer.

TNBC is a highly aggressive subtype of breast cancer characterized by the absence of estrogen receptor (ER), progesterone receptor (PR), and HER2 (ERBB2) expression.^16,17^ The lack of these therapeutic targets limits treatment options, resulting in poor clinical outcomes, early metastasis, and high recurrence rates.^18^ TNBC exhibits greater immunogenicity than other breast cancer subtypes with elevated levels of tumor-infiltrating lymphocytes, PD-L1 expression, and tumor mutational burden.^19,20^ These features make TNBC a promising candidate for immune checkpoint inhibitor (ICI)-based therapies, and the anti-PD1 antibody pembrolizumab is approved for the treatment of TNBC in combination with chemotherapy.^19–21^ Clinical responses to ICIs remain inconsistent, however, and resistance frequently develops, underscoring an incomplete understanding of the regulatory mechanisms governing antitumor immunity in TNBC. Recent single-cell studies have substantially advanced our understanding of the TNBC TME, providing key insights into immune cell composition, subtype signatures, and regulatory functions.^22–30^ Yet, critical gaps persist. In particular, we lack a comprehensive understanding of how circadian regulation organizes the tumor–immune ecosystem in TNBC and how this temporal organization shapes tumor immune evasion. This gap prompts several fundamental questions. How are circadian regulatory networks structured across the TME, spanning both cancer epithelial cells and diverse immune populations? Do cancer epithelial cells harbor intrinsic circadian programs associated with state transitions linked to immune escape? Do cancer and immune compartments exhibit coordinated or divergent circadian programs within individual tumors? How are such temporal relationships associated with functional states of immune activation and suppression? And how might this temporal organization contribute to tumor immune evasion? Addressing these questions will define the temporal architecture of tumor–immune regulation and inform potential chronotherapy strategies.

Addressing these questions has been challenging. Circadian cancer research faces two major barriers: biological phase heterogeneity across individuals and limited temporal sampling of clinical specimens, particularly across the full 24-hour cycle. To enable systematic interrogation of circadian dynamics, we implemented time-series single-nucleus multiomic profiling in the 4T1 murine model of TNBC, allowing reconstruction of circadian regulatory networks linking cancer epithelial and immune compartments within the TME. We uncovered subtype-specific oscillations in cancer epithelial cell states and widespread circadian misalignment between cancer epithelial and immune populations across key functional pathways governing proliferation, antigen presentation, and immune activation. Circadian overlap between T cell activation and exhaustion constrained effective antitumor immunity, while phase mismatch within the PD-1–PD-L1 axis prolonged immune suppression. Together, these findings reveal temporal discoordination as a fundamental feature of tumor–immune interactions in TNBC. To evaluate the translational relevance of these findings, we examined human TNBC datasets. Cross-species comparison revealed comparable circadian gene co-regulation patterns, supporting the potential clinical relevance of circadian misalignment for immunotherapy strategies.

## RESULTS

### Circadian single-cell transcriptomic and epigenomic landscape of breast cancer

To investigate how circadian interactions between immune and non-immune cells modulate immune surveillance and tumor development, we performed single-nucleus multiomic profiling (matched snRNA-seq and snATAC-seq) of 4T1 orthotopic tumors from BALB/cJ mice collected at four circadian time (CT) points (CT4, CT10, CT16, and CT22) (**Figure 1A**). Although tumors collected at nighttime had slightly longer growth periods, the tumor volumes and histopathological features did not differ significantly across time points (**Figures S1A and S1B**). We separated CD45⁺ and CD45⁻ fractions prior to nuclei isolation to improve the capture of both immune-infiltrated and tumor-intrinsic populations, enabling detailed dissection of cell-type-specific circadian programs within the TME. After rigorous quality control and integration of snRNA-seq and snATAC-seq modalities, we obtained 101,321 high-quality nuclei with matched transcriptomic and chromatin accessibility profiles (**Figures 1B and S1C–S1G)**. By integrating snRNA gene expression, snATAC gene activity and transcription factor (TF) motif enrichments, we identified 14 cell types (**Figures 1B–1D**). Within the CD45⁺ compartment, these included CD4⁺ T cells, CD8⁺ T cells, natural killer T (NKT) cells, natural killer (NK) cells, B cells, macrophages/monocytes (Macro/Mono), neutrophils (Neu), dendritic cells (DC), and plasmablasts (**Figures 1B–1D**). Within the CD45⁻ compartment, we identified cancer-associated fibroblasts (CAFs), endothelial cells (Endo), perivascular-like cells (PVLs), normal epithelial cells (Normal Epi), and cancer epithelial cells (Cancer Epi) (**Figures 1B–1D**). Among these cell types, CD4⁺ T cells and cancer epithelial cells were the most abundant within the CD45⁺ and CD45^-^compartments, respectively (**Figure 1B**).

**Figure 1.**
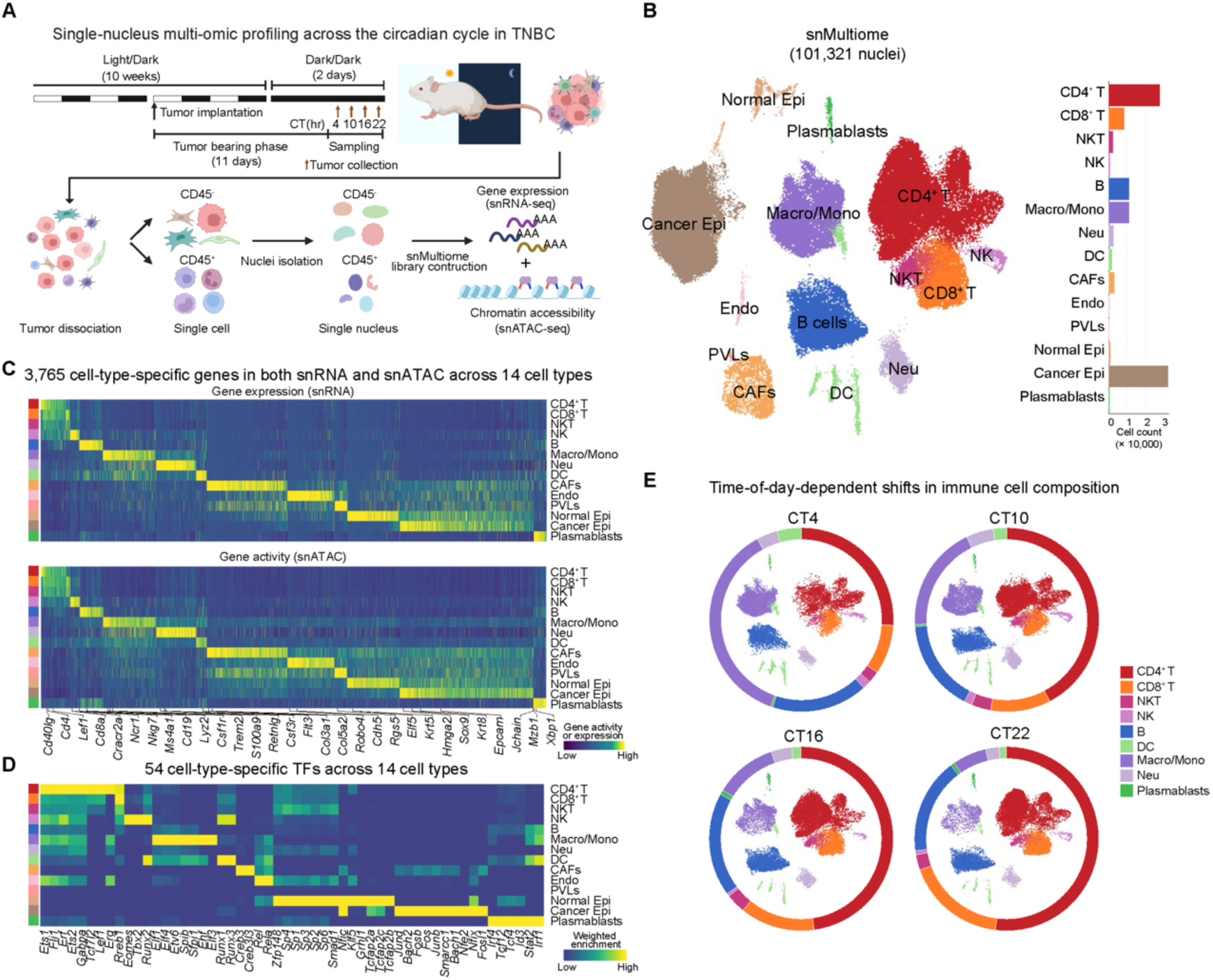
Single-cell epigenomic and transcriptomic landscape of the TNBC tumor microenvironment across circadian time points. (A) Experimental design and analytical workflow. 4T1 tumors were collected from mice maintained under a 12 h light/12 h dark cycle at four circadian time points (CT4, CT10, CT16, CT22) 11 days post-implantation. CD45⁺ and CD45⁻ nuclei were isolated for single-nucleus RNA and ATAC multiomic sequencing (snMultiome). (B) UMAP embedding of 101,321 nuclei profiled by snMultiome, annotated into 14 cell types, including immune (CD4⁺ T, CD8⁺ T, NK, B, NKT, DC, Macro/Mono, Neu, Plasmablast) and non-immune (Cancer Epi, Endo, CAF, PVL, Normal Epi) populations (left). Bar plots depict the relative abundance of each cell type across the full dataset (right). (C) Heatmaps showing cell-type-specific gene expression (snRNA-seq; top) and gene activity inferred from chromatin accessibility (snATAC-seq; bottom) across 14 cell types. (D) Heatmap of 54 cell-type-specific transcription factors across 14 TME cell types, showing motif deviation scores inferred from snATAC-seq. (E) UMAP plots illustrating the temporal distribution of CD45⁺ immune cell types across four circadian time points (CT4, CT10, CT16, and CT22). Stacked ring plots surrounding each UMAP depict the relative proportions of individual immune cell types at each time point, highlighting time-of-day–dependent changes in immune infiltration.

We next examined temporal variation in cell composition and observed pronounced time-of-day–dependent changes in immune infiltration, with CD4⁺ and CD8⁺ T cells peaking at night and Macro/Mono subsets decreasing correspondingly, indicating circadian remodeling of immune composition within the TME (**Figures 1E and S1H**). These data establish a time-resolved single-nucleus multiomic atlas of the tumor microenvironment, enabling dissection of circadian regulation across both cancer and immune cell types.

### Cell-type-specific circadian gene programs reveal functional diversity across the TME

As the relative abundance of tumor-infiltrating cell populations varies across the circadian cycle, we next examined whether these cells also exhibit circadian transcriptional programs. We applied a pseudo-bulk strategy across cell types and circadian time points, followed by DiscoRhythm analysis to identify rhythmic genes (**Figure S2A**).^31^ Temporal dynamics were reconstructed by ordering cells according to circadian sampling time and ranking rhythmic genes by their acrophase, followed by grouping genes into eight phase bins (P1–P8; P1: CT0–3, P2: CT3–6, P3: CT6–9, P4: CT9–12, P5: CT12–15, P6: CT15–18, P7: CT18–21, P8: CT21–24) based on their acrophases (**Figures 2A and S2A; STAR Methods**). In total, we identified 6,224 circadian genes across the eight cell types. Among these, 2,912 were rhythmic in cancer epithelial cells, 2,291 in CD4⁺ T cells, 1,661 in B cells, 1,380 in macrophage/monocyte cells, 786 in CD8⁺ T cells, 721 in CAFs, 459 in neutrophils, and 391 in NKT cells (**Figures 2A and S2A; Table S1**). Across cell types, circadian programs comprised both shared and cell-type-specific components. While only a small subset of genes (e.g., *S100a6*, *Btg1*, *Rps28*) oscillated in all eight cell types, 2,541 genes oscillated in at least two or more cell types, including core clock regulators such as *Arntl*, *Per1*, *Per2*, and *Rora*, reflecting both coordinated and cell-type-specific circadian regulation within the TME (**Figures S2B and S2C**).

**Figure 2.**
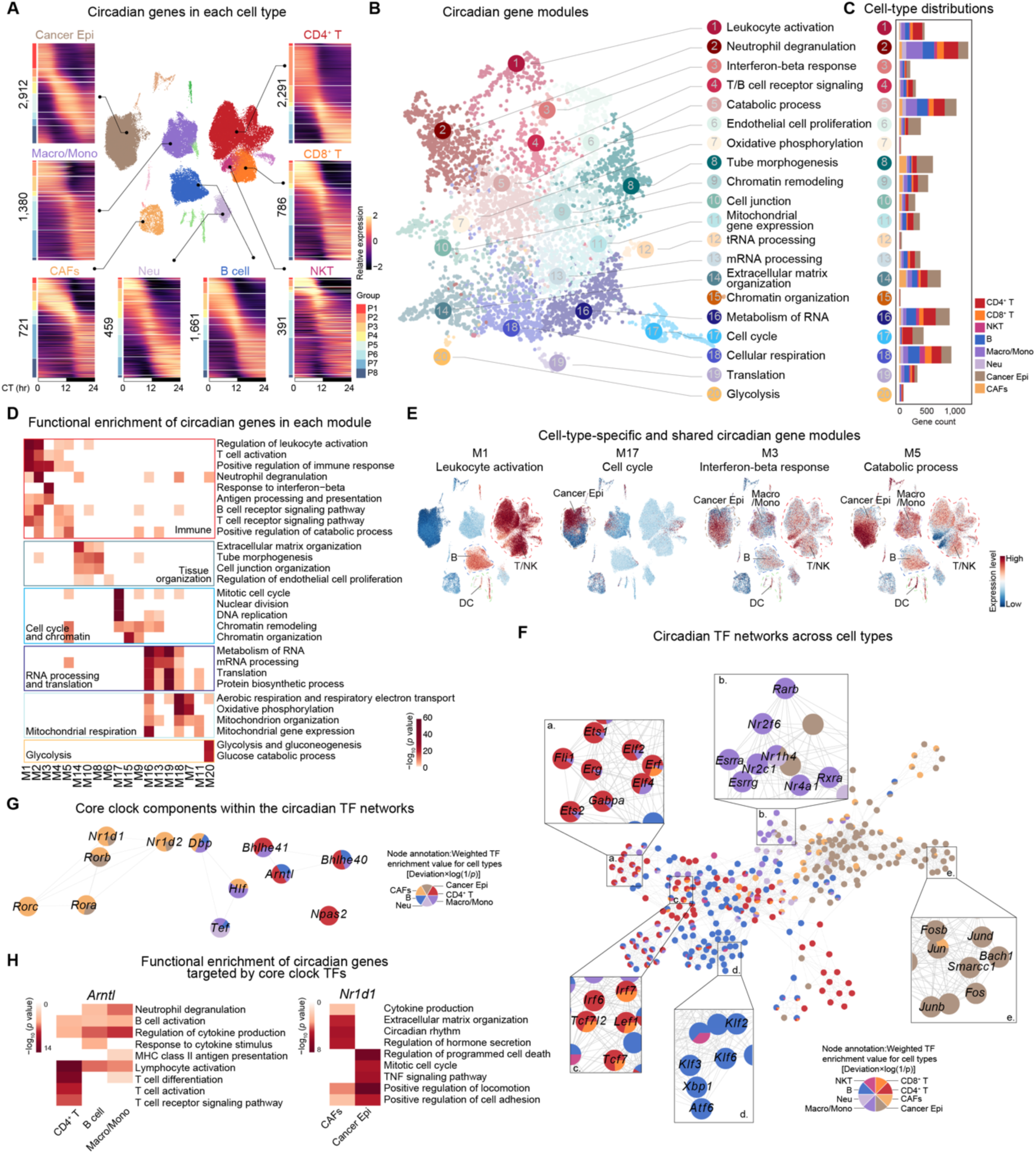
Circadian gene modules and regulatory networks reveal cell-type-specific rhythmic programs across the TME. (A) Heatmap displaying the expression dynamics of circadian genes across cancer epithelial cells (Cancer Epi), cancer-associated fibroblasts (CAFs), macrophage/monocyte (Macro/Mono), neutrophil (Neu), natural killer T (NKT) cells, B cells, CD8⁺ T cells, and CD4⁺ T cells. The number of rhythmic genes identified in each cell type is indicated in black. Heatmap color bar reflects expression level from high (bright yellow) to low (deep purple). (B) Circadian gene modules identified from rhythmic genes across all cell types. Each dot represents a gene, colored by its assigned module, and positioned according to similarity in gene functions. (C) Bar plots showing the relative count of rhythmic genes in each module across TME cell types. (D) Heatmap showing functional enrichment of circadian gene modules. Rhythmic genes in the TME were mainly associated with immune response, tissue organization, cell cycle and chromatin, RNA processing and translation, mitochondrial respiration, and glycolysis. Color intensity reflects the −log_10_(*p* value) of enrichment, ranging from deep red (high) to white (low). (E) UMAP visualizations of selected circadian gene modules. Color intensity reflects the module expression level, ranging from red (high) to blue (low). (F) Circadian TF networks showing motif binding with rhythmic activity across cell types. Each node represents a TF, colored by cell-type enrichment and scaled by the deviation score and weighted rhythmic significance within each cell type. Insets (a–e) highlight representative TF subnetworks enriched in specific cell types (G) Networks representation of core circadian clock TF identified within the circadian TF networks. Nodes represent clock components, colored by cell-type enrichment and scaled by the deviation score and weighted rhythmic significance within each cell type. (H) Heatmaps showing the functional enrichment of circadian genes targeted by core circadian clock TFs *Arntl* (left) and *Nr1d1* (right) across selected TME cell types. The color bar reflects the −log_10_(*p* value) of enrichment, ranging from deep red (high) to white (low).

To systematically characterize the functional architecture of circadian regulation, we constructed gene co-expression modules from the union set of circadian genes identified across cell types and circadian phases. This analysis resolved 20 functional circadian modules representing distinct temporal and biological programs (**Figures 2B and 2C; Table S2; STAR Methods**). These modules encompassed key biological processes, including immune responses (M1–M5), tissue organization (M6, M8, M10, and M14), cell cycle and chromatin (M9, M15, and M17), RNA processing and translation (M13, M16, and M19), mitochondrial respiration (M7, M11, M18), glycolysis (M20), among others (**Figure 2D**). Module gene expression exhibited cell-type specificity. M1 (leukocyte activation) and M4 (T/B cell receptor signaling) circadian genes were higher in CD4⁺ T, CD8⁺ T, and B cells, whereas M2 (neutrophil degranulation) circadian genes were enriched in macrophage/monocyte and neutrophil populations, reflecting circadian immune activation aligned with their cellular functions (**Figures 2E and S2D**). Cancer-specific circadian programs were represented by M8 (tube morphogenesis), M9 (chromatin remodeling) and M17 (cell cycle) (**Figures 2E and S2D**). In addition to cell-type-specific activation of distinct modules, we observed that several circadian programs were shared across cellular compartments. For example, M14 (extracellular matrix organization) was primarily shared between cancer epithelial cells and CAFs, highlighting intercellular coupling across tumor and stromal compartments (**Figure S2D**). Notably, immune signaling modules M3 (interferon-beta response) and M5 (catabolic process) were broadly expressed across both cancer and immune compartments, indicating coordinated circadian regulation of inflammatory and kinase pathways (**Figure 2E**). Similarly, mitochondrial modules M7 (oxidative phosphorylation) and M11 (mitochondrial gene expression) were broadly expressed across multiple cell types (**Figure S2D**). Together, these results demonstrate that circadian regulation in the TME is both cell-type-specific and broadly coordinated across cellular lineages.

We next examined whether genes associated with clinically relevant TNBC therapies were rhythmically regulated. Accordingly, we assessed enrichment of drug-associated gene sets within circadian modules. Chemotherapy target-associated genes were selectively enriched in the cell-cycle module M17, and *Tubb4b*, a β-tubulin isoform targeted by paclitaxel, showed clear oscillations in both cancer epithelial and CD8⁺ T cells, suggesting potential circadian variation in therapeutic efficacy and adverse effects (**Figure S2E)**. In contrast, immune checkpoint blockade target associated genes were enriched in the immune-signaling module M5 (**Figure S2E**). Notably, *Pten*, whose downregulation has been linked to diminished benefit from PD-1/PD-L1 blockade including pembrolizumab,^32,33^ exhibited robust oscillations across cancer epithelial, macrophage/monocyte, and B cell populations, indicating that circadian timing may modulate drug–target availability within the TME (**Figure S2E**).

Collectively, this circadian analysis defines cell-type-specific and shared rhythmic programs across the TME, providing a framework for understanding temporal gene regulation and its potential therapeutic relevance in TNBC.

### Cell-type-specific circadian TF networks

Building on the circadian gene programs identified across cell types, we next sought to identify upstream regulatory drivers. We computed single-cell–resolved TF deviation scores using chromVAR implemented in ArchR, performed pseudo-bulk aggregation across circadian time points for each cell type, and applied DiscoRhythm to identify TFs exhibiting rhythmic regulatory activity (**Figures 2F and S2A; Table S3**). In CD4⁺ T cells, ETS/ELF-family factors (*Erg*, *Ets1*, *Elf2*, *Elf4*) exhibit rhythmic TF activity, supporting rhythmic programs of T cell differentiation and activation (**Figure 2F**).^34^ In both CD4⁺ and CD8⁺ T cells, TCF/LEF factors (*Tcf7*, *Lef1*) and IRF factors (*Irf7*) were implicated in the circadian regulation of T cell differentiation and effector activation (**Figure 2F**).^35–39^ In B cells, KLF factors (*Klf2*, *Klf3*), and *Xbp1* displayed circadian motif activity, linking rhythmic regulation to B-cell activation, metabolism, and differentiation (**Figure 2F**).^40–43^ In Macro/Mono populations, *Nr4a1*, *Esrra*/*Esrrg*, and *Rxra*/*Rarb*, act as circadian TFs, regulating anti-inflammatory programming and mitochondrial and lipid metabolic pathways (**Figure 2F**).^44–46^ In cancer epithelial cells, AP-1 factors (*Fos*, *Fosb*, *Jun*, *Junb*, *Jund*) govern proliferative and stress-responsive transcriptional programs in a circadian manner (**Figure 2F**).^47,48^

Notably, core clock TFs orchestrated circadian regulation in a coordinated yet cell-type-specific manner (**Figure 2G**). *Arntl* governed shared circadian immune programs across CD4⁺ T cells, Macro/Mono, and B cells, including lymphocyte activation and cytokine production, while regulating cell-type-specific targets such as T cell receptor signaling in CD4⁺ T cells, antigen processing in Macro/Mono, and cytokine-response pathways in B cells (**Figures 2G and 2H**). *Nr1d1* regulated adhesion- and locomotion-related pathways in both cancer epithelial cells and CAFs, with lineage-specific control of mitotic cell-cycle genes in cancer epithelial cells and extracellular matrix organization in CAFs (**Figures 2G and 2H**). *Dbp* displayed rhythmic activity across CAFs, B cells, and Macro/Mono, with temporally distinct regulation of immune and migratory pathways (**Figures 2G and S2F**). Importantly, even shared circadian targets exhibited phase divergence across cell types, as exemplified by *Irf7* (regulated by *Arntl*), *Btg1* (*Nr1d1*), and *Hif1a* (*Dbp*), which peaked at distinct circadian phases depending on cellular context (**Figure S2G**).

These findings reveal that circadian transcriptional regulation within the TME is characterized by both cell-type specificity and temporal diversity. Even under shared core clock control, each cell population maintains distinct rhythmic programs and activation phases. These observations highlight that circadian programs can be temporally uncoupled across cellular populations within the TME.

### Circadian misalignment between cancer epithelial and immune cells in the TME

To systematically characterize temporal organization across the TME, we examined the global acrophase distributions of circadian genes across major TME cell types. This analysis revealed an overall phase misalignment between cancer epithelial cells and both stromal and immune compartments (**Figure 3A**). To further define intercellular phase relationships, we compared acrophase offsets of jointly rhythmic genes that exhibited significant rhythmicity in both cell types. These genes were classified as *Aligned* (0–3 h), *Mismatched* (3–8 h), or *Reversed* (8–12 h) based on phase differences (**Figure S3A**). Among immune populations, functionally related cell types exhibited stronger phase concordance, with a large fraction of jointly rhythmic genes classified as *Aligned* (**Figure 3B**). CD8⁺ T cells and NKT cells exhibited the highest concordance, with over 95% of jointly rhythmic genes aligned (237 genes), consistent with their shared effector functions and coordinated circadian regulation of RNA metabolism, oxidative phosphorylation, adaptive immune processes, and cytokine signaling (**Figures 3B and 3C**). Similarly, more than 84% of jointly rhythmic genes between CD4⁺ and CD8⁺ T cells were aligned (397 genes), enriched in pathways of RNA metabolism, T cell activation, lymphocyte proliferation, and T cell differentiation (**Figures 3B and 3D**). These findings indicate that functionally related immune populations maintain synchronized circadian programs within the TME.

**Figure 3.**
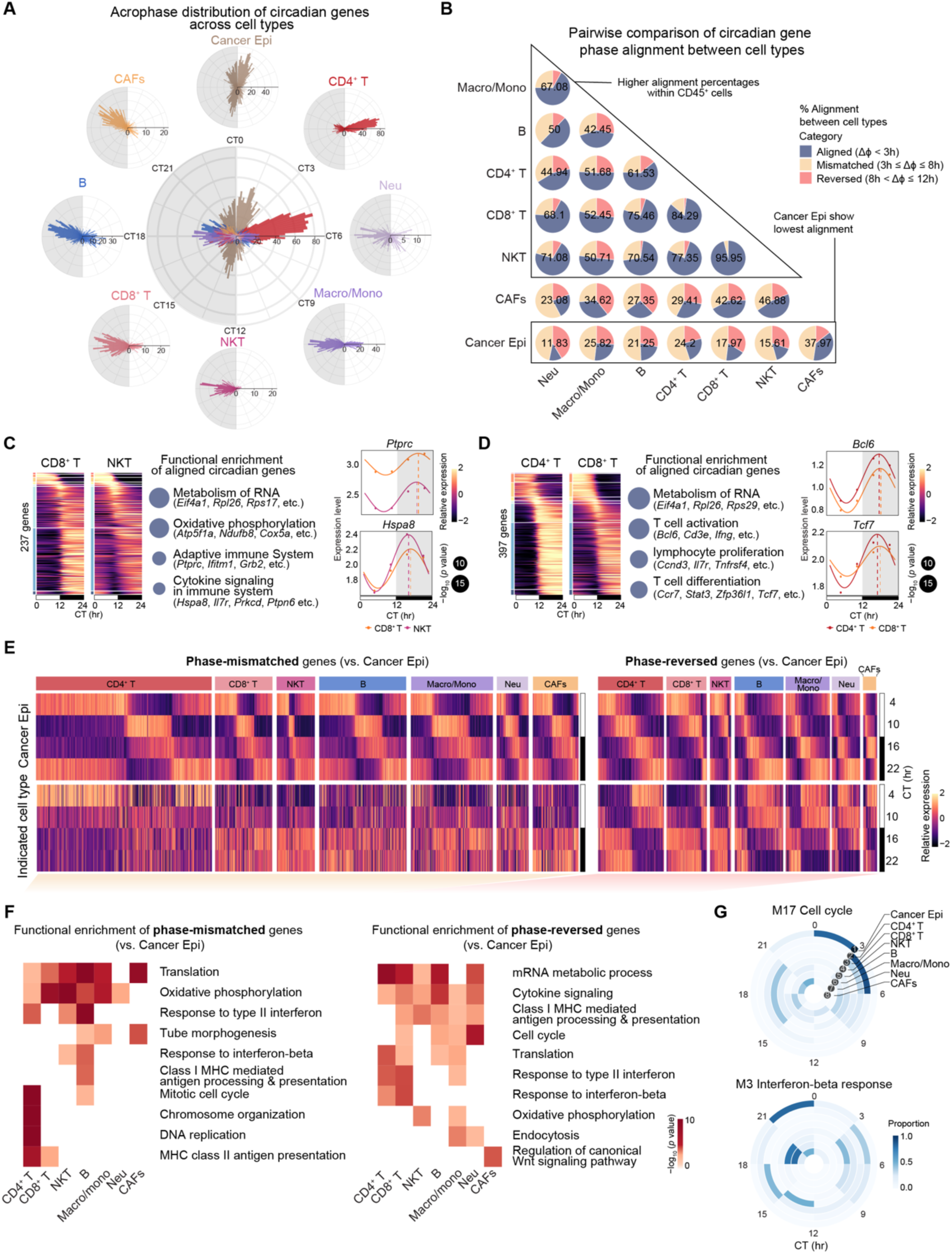
Circadian gene misalignment between cancer cells and other cells in the TME. (A) Radial plots showing the acrophase distribution of circadian genes in each cell type, along with a combined profile across all cell types. The gray area denotes the dark phase (CT12–CT24/CT0). The number of rhythmic genes per group is indicated in black. (B) Pairwise comparison of circadian gene phase alignment between cell types. Pie charts show the proportion of aligned, mismatched, and reversed shared circadian genes for each comparison, with black labels indicating the percentage of aligned genes. Higher alignment percentages are observed among immune cell types, whereas cancer epithelial cells display the lowest alignment. (C) Left: Heatmaps of aligned circadian genes shared by CD8⁺ T and NKT. Heatmap color bar reflects expression level from high (bright yellow) to low (deep purple). Middle: Dot-plot of functional enrichment of these aligned genes. Dot size encodes −log_10_(*p* value). Right: Circadian fitting curves showing the expression dynamics of *Ptprc* and *Hspa8* across a circadian day in CD8⁺ T and NKT cells. (D) Left: Heatmaps of aligned circadian genes shared by CD4⁺ and CD8⁺ T cells. Heatmap color bar reflects expression level from high (bright yellow) to low (deep purple). Middle: Dot-plot of functional enrichment of these aligned genes. Dot size encodes −log_10_(*p* value). Right: Circadian fitting curves showing the expression dynamics of *Bcl6* and *Tcf7* across a circadian day in CD4⁺ and CD8⁺ T cells. (E) Heatmaps showing the expression patterns of circadian genes that are phase-mismatched (left) and reversed (right) between cancer epithelial cells and other cell types. The color bar reflects the expression level from high (bright yellow) to low (deep purple). (F) Heatmaps showing the functional enrichment of circadian genes that are phase-mismatched (left) and reversed (right) between cancer epithelial cells and other cell types. The color bar reflects the −log_10_(*p* value) of enrichment, ranging from high (red) to low (white). (G) Circular plots showing the acrophase distribution of genes within M17 (top) and M3 (bottom) circadian modules across different cell types. The color bar indicates the proportion of circadian genes peaking within each of the eight phase bins (P1–P8), from low (white) to high (dark blue).

In contrast to the coordinated rhythmicity observed among immune populations, cancer epithelial cells exhibited widespread phase misalignment with stromal and immune compartments (**Figures 3A, 3B, 3E and S3B**). Phase-mismatched genes between cancer epithelial and immune cells were enriched in translation, oxidative phosphorylation, and interferon response-related pathways across multiple comparisons (**Figure 3F**). Genes mismatched with CD4⁺ T cells additionally showed enrichment in cell-cycle and DNA replication programs, whereas mismatched genes involving CD4⁺ T, CD8⁺ T, or B cells were associated with antigen processing and presentation (**Figure 3F**). Phase-reversed genes were enriched in RNA metabolism and cytokine signaling pathways across several immune cell types (**Figure 3F**). Notably, antigen presentation and interferon-response programs were enriched in comparisons involving CD4⁺ T cells, CD8⁺ T cells, and macrophage/monocyte populations (**Figure 3F**). At the module level, circadian modules linked to cell cycle (M17), interferon-β response (M3), metabolism of RNA (M16), translation (M19), and oxidative phosphorylation (M7) similarly exhibited phase divergence relative to cancer epithelial cells (**Figures 3G and S3C**).

Together, these findings reveal a cell-type-specific circadian organization within the TME, characterized by coordinated rhythmic programs among immune populations but widespread temporal misalignment between cancer epithelial and immune compartments, resulting in temporal decoupling of proliferative and immune regulatory programs within the TME.

### Circadian state switching in cancer epithelial cells

Given the observed phase misalignment between cancer and immune cells, we next investigated the circadian regulation in cancer epithelial cells. Single-nucleus CNV profiling confirmed their malignant identity (**Figure S4A**). Phase-resolved analysis of 2,912 circadian genes revealed a clear temporal ordering of transcriptional programs across the circadian cycle, with peaks shifting from proliferation and metabolic activity to antigen presentation and interferon responses (**Figure 4A**). Genes associated with cell cycle, DNA repair, and DNA replication reached peak expression around the late-night to early-day transition (CT23-3), whereas antigen-presentation and interferon-response genes peaked during the late night (CT20–24) (**Figures 4A, 4B, and S4B–S4G**). Gene signature scores confirmed a time-of-day–dependent alternation between proliferative programs dominating around the late-night to early-day transition and immunogenic activity peaking during the late night (**Figure S4H**).

**Figure 4.**
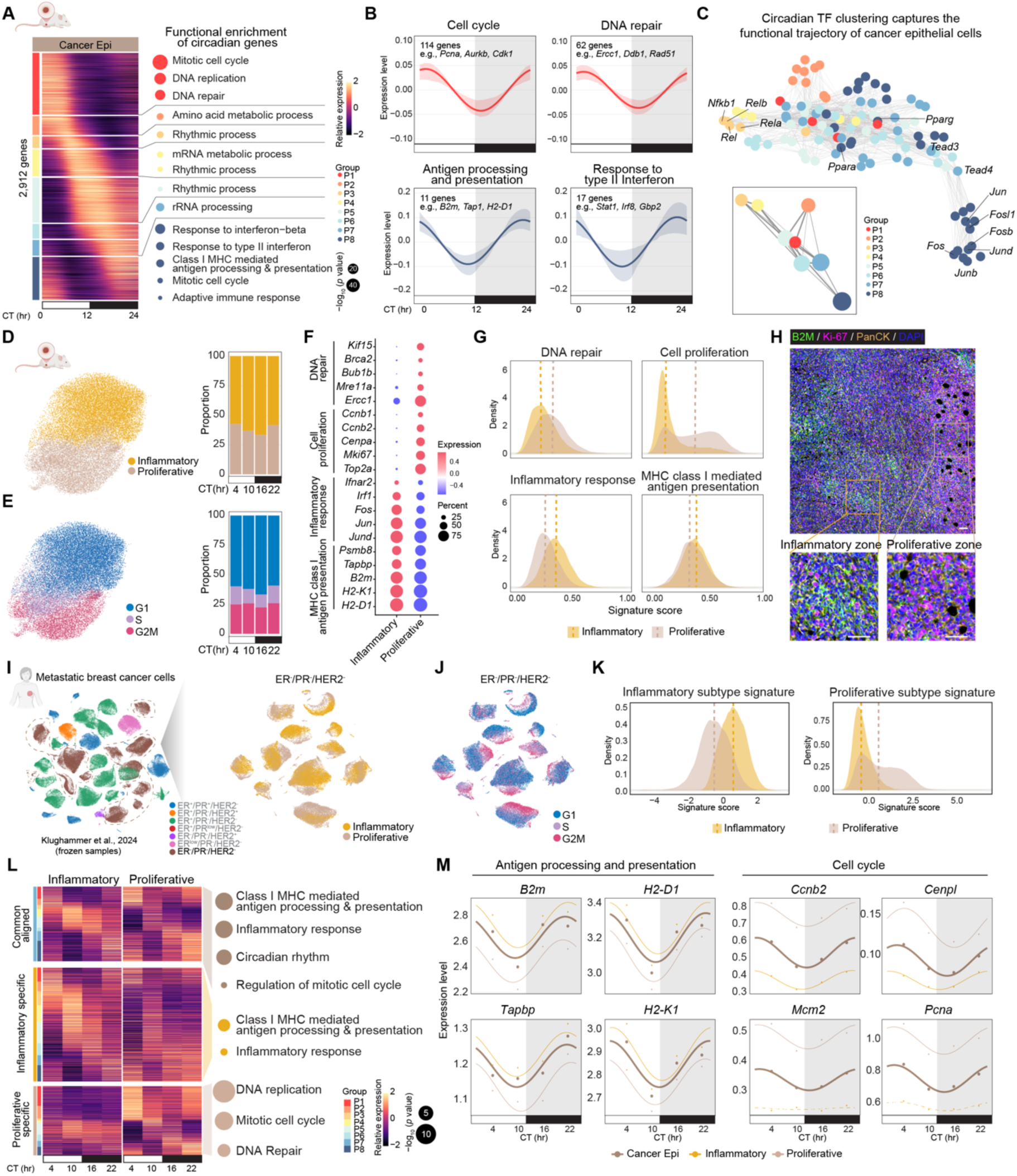
Time-of-day–driven oscillation between proliferative and inflammatory states in cancer epithelial cells. (A) Heatmap showing rhythmic gene expression in cancer epithelial cells across circadian phases, with corresponding functional enrichment (right). The color bar represents expression level from high (bright yellow) to low (deep purple). Dot size encodes the −log_10_(*p* value) of enrichment. (B) Circadian fitting curves showing the expression dynamics of indicated pathways (cell cycle, DNA repair, antigen processing and presentation, and response to type II interferon) in cancer epithelial cells across a day. Bold lines represent the mean fitted circadian curve across all genes involved in each pathway, and shaded areas indicate the interquartile range (25th–75th percentile). The number of genes included in each pathway is indicated within each panel, along with representative genes. The gray area indicates the dark phase (CT12–CT24/CT0). (C) UMAP and Partition-based graph abstraction (PAGA, bottom left inset) representations illustrating the functional connectivity of circadian transcription factors (TFs) in cancer epithelial cells across circadian phases. Each node represents a TF, colored by its peak phase (P1–P8). TFs associated with cell proliferation and immune response are labeled. The inset (bottom left) shows a low-dimensional representation of phase-specific TF organization, highlighting the sequential transition of TF activity across a day. (D) Left: UMAP visualization showing the distribution of two cancer epithelial subtypes, inflammatory and proliferative cancer epithelial cells. Right: Stacked bar plots displaying the proportion of inflammatory and proliferative cancer epithelial cells. across four circadian time points (CT4, CT10, CT16, and CT22). (E) Left: UMAP visualization of cancer epithelial cells colored by cell-cycle phases (G1, S, and G2/M). Right: Stacked bar plots showing the proportion of cells in each cell-cycle phase across circadian time points (CT4, CT10, CT16, and CT22), revealing rhythmic variation in cycling activity. (F) Dot plot showing expression levels of canonical DNA repair, cell proliferation, inflammatory response and MHC class I mediated antigen presentation markers in inflammatory and proliferative cancer epithelial cells. Dot size indicates the percentage of expressing cells, and color reflects average expression level from high (red) to low (blue). (G) Density plots showing the distribution of functional signature scores, including DNA repair, cell proliferation, inflammatory response, and MHC class I mediated antigen presentation, across inflammatory and proliferative cancer epithelial cells. (H) Representative immunofluorescence staining showing distinct spatial localization of proliferative (PanCK⁺, Ki-67⁺) and inflammatory (PanCK⁺, B2M^high^) cancer epithelial cells within tumor tissues. Scale bars, 50 μm. (I) UMAP projections of single-nucleus RNA-seq data from metastatic breast cancer (MBC) epithelial cells.^49^ Left, cells annotated by molecular subtype. Right, ER^−^/PR^−^/HER2^−^ MBC cells subset from the left panel, annotated by proliferative and inflammatory states. (J) UMAP visualizations of single-nucleus RNA-seq data from ER^−^/PR^−^/HER2^−^ MBC cells subset,^49^ annotated by cell-cycle phase. (K) Density plots showing the distribution of inflammatory and proliferative state signature scores in human ER^−^/PR^−^/HER2^−^ MBC cells,^49^ confirming the presence of analogous states in patients. (L) Heatmap showing rhythmic gene expression patterns in inflammatory and proliferative cancer epithelial cells across circadian time points. Genes are categorized into common aligned, and subtype-specific rhythmic groups, with corresponding functional enrichment shown on the right. The color bar reflects expression level from high (bright yellow) to low (deep purple). Dot size indicates the −log_10_(*p* value) of pathway enrichment, and dot color represents the cell type category (common, inflammatory or proliferative). (M) Circadian fitting curves showing the expression dynamics of representative genes across a circadian day in Cancer Epi (brown bold line), proliferative (gray), and inflammatory (yellow) subtypes. Dots represent the mean expression values at each sampled time point in Cancer Epi. Dashed lines indicate non-significant rhythmicity in that cell type. The gray area indicates the dark phase (CT12–CT24/CT0).

Transcription factors regulating these programs displayed a parallel temporal organization, as reflected by rhythmic motif accessibility. Motifs corresponding to AP-1, TEAD, and PPAR family members (*Fos*, *Jun*, *Tead3/4*, *Ppara*, *Pparg*) peaked during the late-night phase (P8), associated with proliferative and metabolic programs, whereas motifs corresponding to NF-κB family factors (*Nfkb1*, *Rel*, *Rela*, *Relb*) peaked at phase P3, aligned with immune-regulatory activity (**Figure 4C**). Consistent with the expected temporal delay between epigenomic TF activity and downstream gene expression, motif accessibility and target gene expression exhibited offset yet sequential phase patterns, suggesting temporally ordered regulation linking TF activity to proliferative, metabolic, and immune-associated programs in cancer epithelial cells (**Figure 4C**).

### Subtype composition and intrinsic rhythmicity jointly shape cancer epithelial circadian programs

We next asked whether these circadian programs reflect changes in the relative abundance of epithelial subtypes or intrinsic rhythmic regulation within each state. Based on cell-cycle scores and transcriptional signatures, we identified two major cancer epithelial cell states: a proliferative state and an inflammatory state (**Figures 4D and 4E**). Proliferative cancer epithelial cells were predominantly in the G2/M and S phases and exhibited higher expression of cell proliferation, DNA repair, and DNA replication genes, including *Mki67, Ccnb1, Cenpa, Top2a, Brca2, Bub1b,* and *Mre11a* (**Figures 4E–4G and S5A and S5B**). In contrast, inflammatory cancer epithelial cells were enriched in the G1 phase and displayed higher expression of genes involved in antigen processing and presentation (e.g., *B2m, Psmb8, Tapbp, H2-K1*, and *H2-D1*) and inflammatory signaling (e.g., *Irf1* and *Ifnar2*), suggesting an immunomodulatory phenotype (**Figures 4E–4G and S5A**). Consistent with this observation, signature scoring based on circadian module genes revealed that proliferative cancer epithelial cells had higher scores for the cell cycle associated module (M17), whereas inflammatory cancer epithelial cells had higher scores for the interferon-β response module (M3) (**Figure S5C**). These transcriptional differences were further supported at the protein level by immunofluorescence, which revealed Ki-67 positive cancer epithelial cells with lower B2M expression and Ki-67 negative cancer epithelial cells with higher B2M abundance (**Figures 4H and S5D**). Single-nucleus CNV analysis revealed comparable chromosomal alteration patterns between the two groups, indicating that these states reflect transcriptional variation within the same malignant clone rather than genetically distinct subpopulations (**Figure S5E**). The proportions of these states oscillated across the circadian cycle, with inflammatory cells peaking around CT16, indicating a temporal transition between proliferative and inflammatory states in cancer epithelial cells (**Figures 4D and 4E**). Moreover, these two states were conserved in human TNBC tumors. Analysis of scRNA-seq datasets revealed that human cancer epithelial cells could also be separated into proliferative and inflammatory populations based on cell-cycle phase, mirroring the proliferative and inflammatory signatures observed in mouse tumors (**Figures 4I–4K and S5F and S5G**).^28,49^

Consistent with their transcriptional signatures, the two cell states exhibited distinct circadian programs. While both subtypes shared circadian regulation of genes involved in rhythmic processes, inflammatory responses, MHC class I antigen presentation (*B2m*, *H2-D1*, *Tapbp*, *H2-K1*), and cell-cycle (*Ccnb2*, *Cenpl*), inflammatory cancer epithelial cells exhibited higher overall expression levels and stronger rhythmic amplitudes of MHC class I antigen-processing genes. (**Figures 4L, 4M, and S5H**). By contrast, proliferative cancer epithelial cells exhibited subtype-specific rhythmic activation of mitotic cell-cycle, DNA replication, and DNA repair genes, including *Mcm2* and *Pcna*, mainly peaking at phase P1 (CT0–3) (**Figures 4L, 4M and S5I**). Circadian TF analysis revealed both shared and subtype-specific regulatory programs that mirror the distinct functional states. A core set of AP-1 family members (*Fos* and *Jun*) together with oxidative and redox-responsive regulators (*Bach1*/*Bach2* and *Nfe2l2*) formed a circadian stress-adaptation module that integrates metabolic and redox homeostasis, representing a common regulatory backbone across subtypes (**Figure S5J**). Building on this shared regulatory TF architecture, inflammatory cells engaged NF-κB pathway TFs (*Rela, Nfkb1*) and C/EBP family members (*Cebpb, Cebpd*), driving circadian activation of antigen-presentation and inflammatory programs, whereas proliferative cells selectively enriched nuclear receptor–based circadian regulators (*Rxra, Nr2f2*) to sustain rhythmic transcriptional control of proliferation (**Figure S5J**).

Together, these findings demonstrate that circadian regulation within cancer epithelial cells emerges from the interplay between time-varying subtype composition and cell-intrinsic, subtype-specific rhythmic programs, producing a robust, time-of-day–dependent shift between proliferative and inflammatory states.

### Coordinated circadian activation of T cells and Macro/Mono at night

Having defined the circadian dynamics of cancer epithelial cells, we next examined the temporal organization of immune populations within the tumor microenvironment. Our analysis revealed that the proportion of functional T and NK cell subtypes peaked during the nighttime (CT16), corresponding to the murine active phase (**Figures S6A–S6E**). Across the circadian cycle, CD4⁺ and CD8⁺ T cells exhibited a temporal transition from cell-cycle and immune-inhibitory states during the early daytime to immune-activation and interferon-response programs at night (**Figures 5A and 5B**). Signature score analysis further revealed timepoint-dependent change of T cell functional states, with effector and interferon-response activities elevated at night, whereas anergy and exhaustion signatures predominated during the day (**Figure 5C**). Together, these findings demonstrate that T cell activation and effector functions follow a robust circadian rhythm, aligning with the mouse’s active (nighttime) phase.

**Figure 5.**
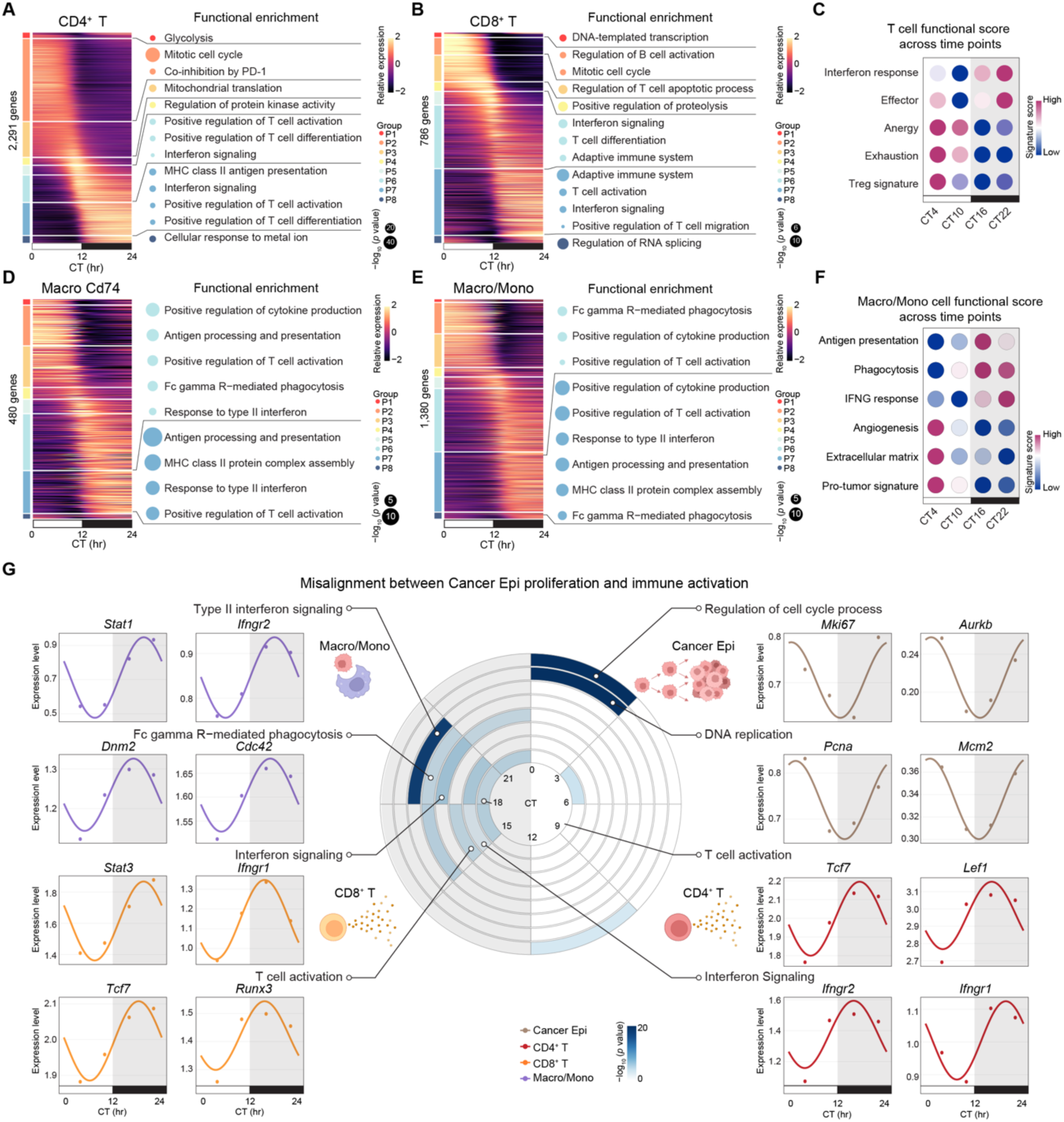
Temporal misalignment of Cancer Epi proliferation and immune activation. (A-B) Heatmap showing rhythmic expression patterns of circadian genes in CD4⁺ T (A) and CD8⁺ T (B) cells across circadian phases. The color bar reflects the expression level from high (bright yellow) to low (deep purple). The adjacent bubble plot displays enriched biological processes peak in each circadian phase group. Dot size indicates the −log_10_(*p* value) of enrichment, and dot color denotes the corresponding phase group. (C) Dot plot showing the circadian dynamics of T cell functional signature scores across circadian time points. The color bar represents the average signature score, ranging from high (magenta) to low (blue). (D-E) Heatmap showing rhythmic expression patterns of circadian genes in Macro Cd74 subtype (D) and Macro/Mono (E) cells across circadian phases. The color bar reflects the expression level from high (bright yellow) to low (deep purple). The adjacent bubble plot displays enriched biological processes peak in each circadian phase group. Dot size indicates the −log_10_(*p* value) of enrichment, and dot color denotes the corresponding phase group. (F) Dot plot showing the circadian dynamics of Macro/Mono cell functional signature scores across circadian time points. The color bar represents the average signature score, ranging from high (magenta) to low (blue). (G) Polar plot showing functional enrichment of the indicated pathways across cell types (Cancer Epi, CD4⁺ T, CD8⁺ T, and Macro/Mono) across the eight circadian phases. The color bar represents −log_10_(*p* value) from high (dark blue) to low (white). Line plots display the circadian fitting curves of representative genes involved in cell cycle and DNA replication (Cancer Epi), T cell activation (CD4⁺ T and CD8⁺ T), interferon associated signaling (Macro/Mono, CD4⁺ T, CD8⁺ T), and phagocytosis (Macro/Mono). Dots represent the mean expression values at each sampled time point. The gray area indicates the dark phase (CT12–CT24/CT0).

We next analyzed the circadian dynamics of macrophage and monocyte populations within the TME. Sub-clustering identified distinct macrophage subtypes, including a CD74⁺ macrophage population characterized by a higher pro-inflammatory macrophage score, and enhanced antigen-presentation activity (**Figures S6F and S6G**). Compared with other macrophage subtypes, Cd74⁺ macrophages governed a greater number of subtype-specific circadian genes enriched for antigen processing and presentation, response to type II interferon, and phagocytosis, suggesting a dominant anti-tumor role of this subset within the TME (**Figure S6H**). Phase-resolved analysis showed that genes involved in Fc-gamma R-mediated phagocytosis and type II interferon signaling peaked during the nighttime (**Figure 5D**). A similar temporal pattern was observed when analyzing macrophage/monocyte populations, where both circadian pathway analysis and overall signature scoring revealed that phagocytosis, antigen presentation, and interferon-response programs peaked at night (**Figures 5E and 5F**). Together, these findings reveal that both T cells and macrophages exhibit coordinated circadian activation of anti-tumor immune programs, reinforcing the alignment of immune effector activity with the organism’s active phase.

### Circadian misalignment disrupts temporal coordination of anti-tumor immune responses

We next compared circadian phase patterns between cancer epithelial proliferative programs and immune activation pathways within the TME. Cell cycle (*Mki67, Aurkb*) and DNA replication genes (*Pcna, Mcm2*) in cancer epithelial cells peaked during the early-day phase (P1; CT0–3), whereas key immune activation pathways, including T cell activation (*Tcf7, Lef1, Runx3*), interferon signaling (*Ifngr1, Ifngr2, Stat3*) in CD4⁺ or CD8⁺ T cells, and Fc gamma R–mediated phagocytosis (*Dnm2, Cdc42*) and type II interferon signaling (*Stat1, Ifngr2*) in Macro/Mono, peaked during the nighttime (**Figures 5G and S6I–S6K**). This antiphase relationship reflects a circadian decoupling between tumor proliferation and immune activation.

We then analyzed the temporal coordination between tumor antigen presentation and T cell activation. Antigen presentation genes in cancer epithelial (*Tap1, B2m*) and macrophage/monocyte cells (*H2-Eb1, H2-Aa*) peaked at night, whereas T cell recognition genes in CD4⁺ (*Cd3e, Lck*) and CD8⁺ T cells (*Cd3e, Trbc1*) reached their peak phases during daytime (**Figures 6A, S4E, and S7A**). Protein-level analyses recapitulated the transcriptomic misalignment, with B2M protein in PanCK⁺ tumor epithelial cells peaking during CT22–CT4, while CD3⁺ cell abundance was highest at CT10–16 (**Figures 6B and 6C**). These findings indicate that antigen availability and T cell responsiveness are temporally out of phase at both the transcript and protein levels within the TME. In addition to this intercellular phase discordance, we observed temporal overlap of T cell recognition and proliferative programs with exhaustion signatures within T cells (**Figure 6D**). In CD4⁺ T cells, genes involved in T cell recognition (*Cd3e, Lck*), costimulation (*Cd28, Icos, Tnfsf14*), and cell cycle (*Cdk6, Mcm6, Aurkb*) peaked at the same circadian phase as immune checkpoint and exhaustion markers (*Ctla4, Tox*), indicating a synchronous rise of activation and inhibitory circuits (**Figures 6A, 6D–6F and S7B**). Similarly, in CD8⁺ T cells, although *Ctla4* and *Tox* did not exhibit significant rhythmicity, both markers showed higher expression during the daytime, coinciding with peaks in T cell recognition (*Cd3e, Trbc1*) and cell-cycle genes (*Ccnd3, Cks2, Tuba1a*) (**Figures 6A, 6D–6F and S7C**). In addition, exhaustion scores were highest during the daytime phases (CT4–CT10), aligning with the peak phases of T cell recognition and costimulation, reflecting a temporal convergence between activation and inhibitory programs (**Figure 6G**). At the cellular level, proliferating CD4⁺ T cells at CT4 were significantly enriched for the exhausted phenotype (CD4⁺ T_EX_), suggesting that heightened activation during this phase may predispose T cells to functional exhaustion (**Figures S7D and S7E**). Together, these results reveal that circadian phase misalignment shapes the temporal architecture of immune recognition at both intercellular and intrinsic levels, characterized by discordant antigen presentation and T cell recognition and by temporal convergence of activation and exhaustion programs within T cells.

**Figure 6.**
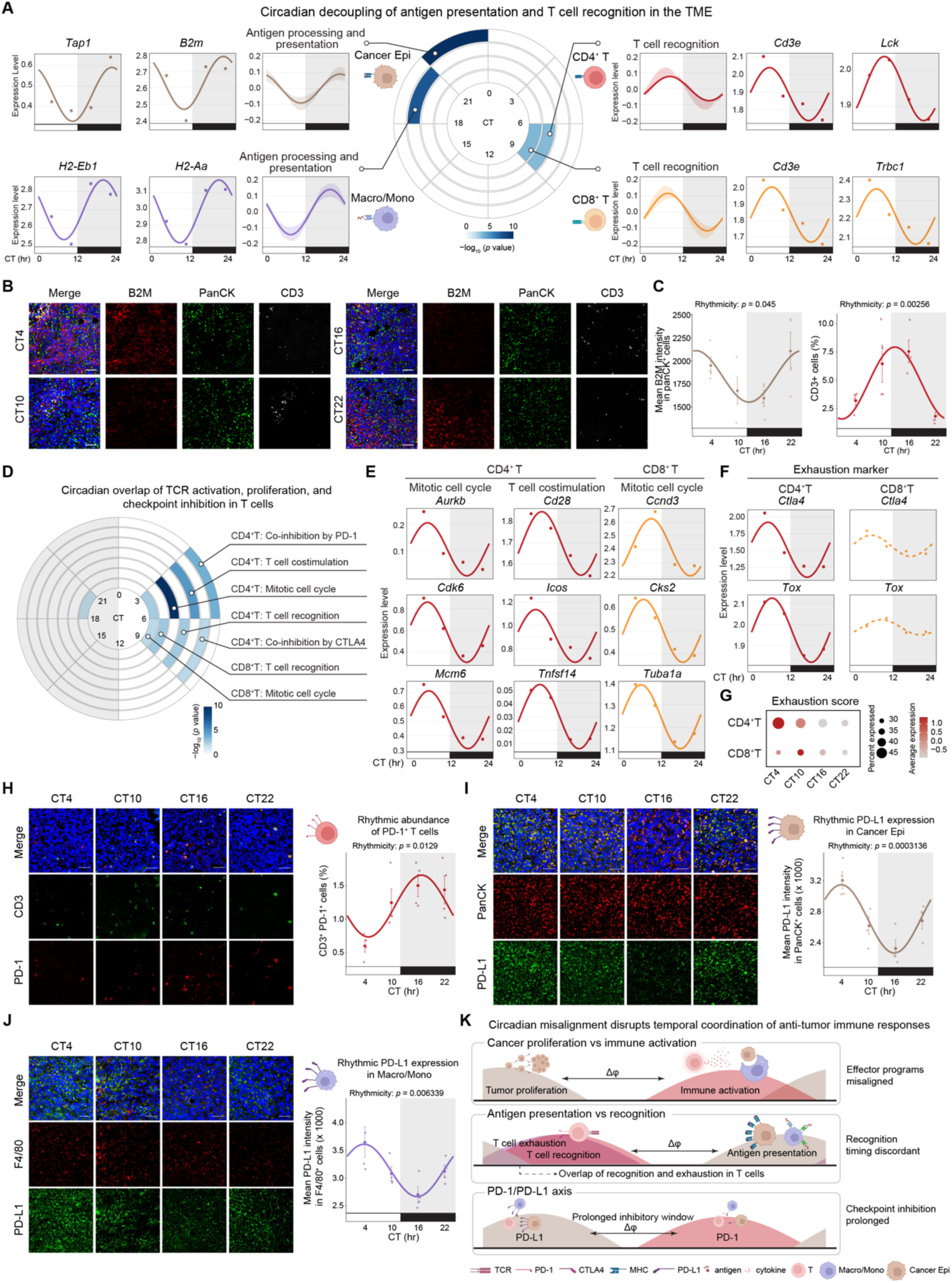
Temporal misalignment between antigen presentation and recognition in TME. (A) Polar plot showing functional enrichment of the indicated pathways involved in antigen presentation (Cancer Epi and Macro/Mono) and T cell recognition (CD4⁺ and CD8⁺ T cells) across the eight circadian phases. The color bar represents −log_10_(*p* value) from high (dark blue) to low (white). Circadian fitting curves showing the expression dynamics of these pathways across a circadian day, with shaded areas indicating the interquartile range (25th–75th percentile). Line plots showing circadian fitting curves of representative genes from each pathway, and dots represent mean expression values at each sampled time point. The gray area indicates the dark phase (CT12–CT24/CT0). (B) Representative immunofluorescence images of tumor sections collected at the indicated circadian times (CT), showing B2M (red), PanCK (green), CD3 (white), and DAPI (blue). Scale bars, 50 μm. (C) Quantification of mean B2M fluorescence intensity in PanCK⁺ tumor epithelial cells (left) and the percentage of CD3⁺ cells (right) across circadian time. Lines indicate cosinor fits, and shaded regions denote the dark phase (CT12–CT24/CT0). *p* values indicate rhythmicity assessed by cosinor regression. *n*=4 tumors per time point. (D) Polar plot showing phase-specific enrichment of pathways associated with T cell recognition, costimulation, proliferation, and checkpoint inhibition across CD4⁺ and CD8⁺ T cells. The color bar represents −log_10_(*p* value) from high (dark blue) to low (white). (E) Line plots showing circadian fitting curves of representative genes involved in mitotic cell cycle and T cell costimulation in CD4⁺ or CD8⁺ T cells. Dots represent mean expression values at each circadian time point. The gray area indicates the dark phase (CT12–CT24/CT0). (F) Line plots showing circadian fitting curves of representative exhaustion markers (*Ctla4* and *Tox*) in CD4⁺ and CD8⁺ T cells. Dots represent mean expression values at each circadian time point. The gray area indicates the dark phase (CT12–CT24/CT0). Dashed lines indicate non-significant rhythmicity in that cell type. (G) Bubble plot showing T cell exhaustion signature scores across circadian time points in CD4⁺ and CD8⁺ T cells. Dot size indicates the percentage of expressing cells, and color reflects average expression levels from high (magenta) to low (blue). (H) Representative immunofluorescence images of tumor sections collected at the indicated circadian times (CT), showing CD3 (green), PD-1 (red), and DAPI (blue). Scale bars, 50 μm. Right, quantification of the percentage of CD3⁺PD-1⁺ T cells among total cells across a circadian day. Lines indicate cosinor fits, and shaded regions denote the dark phase (CT12–CT24/CT0). *p* values indicate rhythmicity assessed by cosinor regression (24-h period). *n* = 4 tumors per time point. (I–J) Representative immunofluorescence images of tumor sections collected at the indicated circadian times, showing PD-L1 (green) in PanCK⁺ Cancer Epi (red; I) or F4/80⁺ Macro/Mono populations (red; J), with DAPI (blue). Scale bars, 50 μm. Right, quantification of mean PD-L1 fluorescence intensity in PanCK⁺ Cancer Epi (I) and F4/80⁺ Macro/Mono populations (J) across a circadian day. Lines indicate cosinor fits, and shaded regions denote the dark phase (CT12–CT24/CT0). *p* values indicate rhythmicity assessed by cosinor regression. *n* = 4 tumors per time point. (K) Schematic model of hierarchical circadian phase offsets across tumor–immune interactions in the TME.

Importantly, circadian phase misalignment extends to the PD-1/PD-L1 immune checkpoint axis, a central determinant of T cell inhibitory signaling and a primary target of immunotherapy.^50–52^ We found that *Pdcd1* expression in CD4⁺ T cells peaked at CT4, consistent with increased chromatin accessibility at the *Pdcd1* locus observed in both CD4⁺ and CD8⁺ T cells during this phase (**Figures S7F and S7G**). In contrast, *Cd274* (encoding PD-L1) expression in both cancer epithelial cells and Macro/Mono reached its maximum much later, around CT22 (**Figure S7H**). Mirroring the transcriptomic misalignment between *Pdcd1* and *Cd274*, this temporal offset was preserved at the protein level, despite a phase shift of protein expression relative to mRNA. PD-1 expression in T cells peaked during the early nighttime, concordant with previously reported circadian regulation of PD-1 protein in T cells,^53^ whereas PD-L1 expression in both cancer epithelial cells and Macro/Mono peaked during the early daytime (**Figures 6H–6J**). This temporal mismatch suggests that phase-offset expression of PD-1 in T cells and PD-L1 in tumor and myeloid cells may prolong checkpoint-mediated inhibitory signaling across the circadian cycle.

Together, these findings reveal a hierarchical pattern of circadian misalignment within the tumor–immune ecosystem, including (1) decoupling between cancer cell proliferation and immune activation, (2) discordant antigen presentation and T cell recognition with intrinsic overlap of recognition/proliferation and exhaustion programs within T cells, and (3) asynchronous PD-1/PD-L1 rhythms across tumor and immune compartments (**Figure 6K**). Collectively, these mechanisms establish a time-of-day–dependent framework suggesting that circadian desynchronization may undermine coordinated anti-tumor immune responses and favor immune evasion within the TME.

### Parallel temporal organization of immune regulation across mouse and human tumors

To assess whether the circadian immune dynamics identified in mouse TME are shared in humans, we inferred the circadian phase in human TNBC samples using the autoencoder CYCLOPS 2.0.^54,55^ Given the cell-type-specific circadian regulation observed in the mouse TME, we constructed an immune-informed circadian seed gene set by combining circadian genes from mouse CD4⁺ T cells, CD8⁺ T cells, and Macro/Mono populations, and apply this framework to a single-cell transcriptomic dataset comprising 76 human TNBC samples (**Figures 7A and S8A**).^23^ After ordering, we found that circadian phase relationships across immune cell types remained highly concordant (**Figure S8B**). Inferred patient-level circadian phases were non-uniformly distributed, with enrichment in specific circadian intervals, consistent with expected clinical sampling bias (**Figure S8C**). Interestingly, we observed circadian variation in immune cell composition within the TME, with T cells enriched at inferred daytime phases and myeloid cells preferentially enriched at inferred nighttime phases (**Figure 7B**). This pattern parallels the temporal organization observed in mouse tumors, while exhibiting the expected phase shift consistent with inverted immune oscillations between diurnal humans and nocturnal mice.

**Figure 7.**
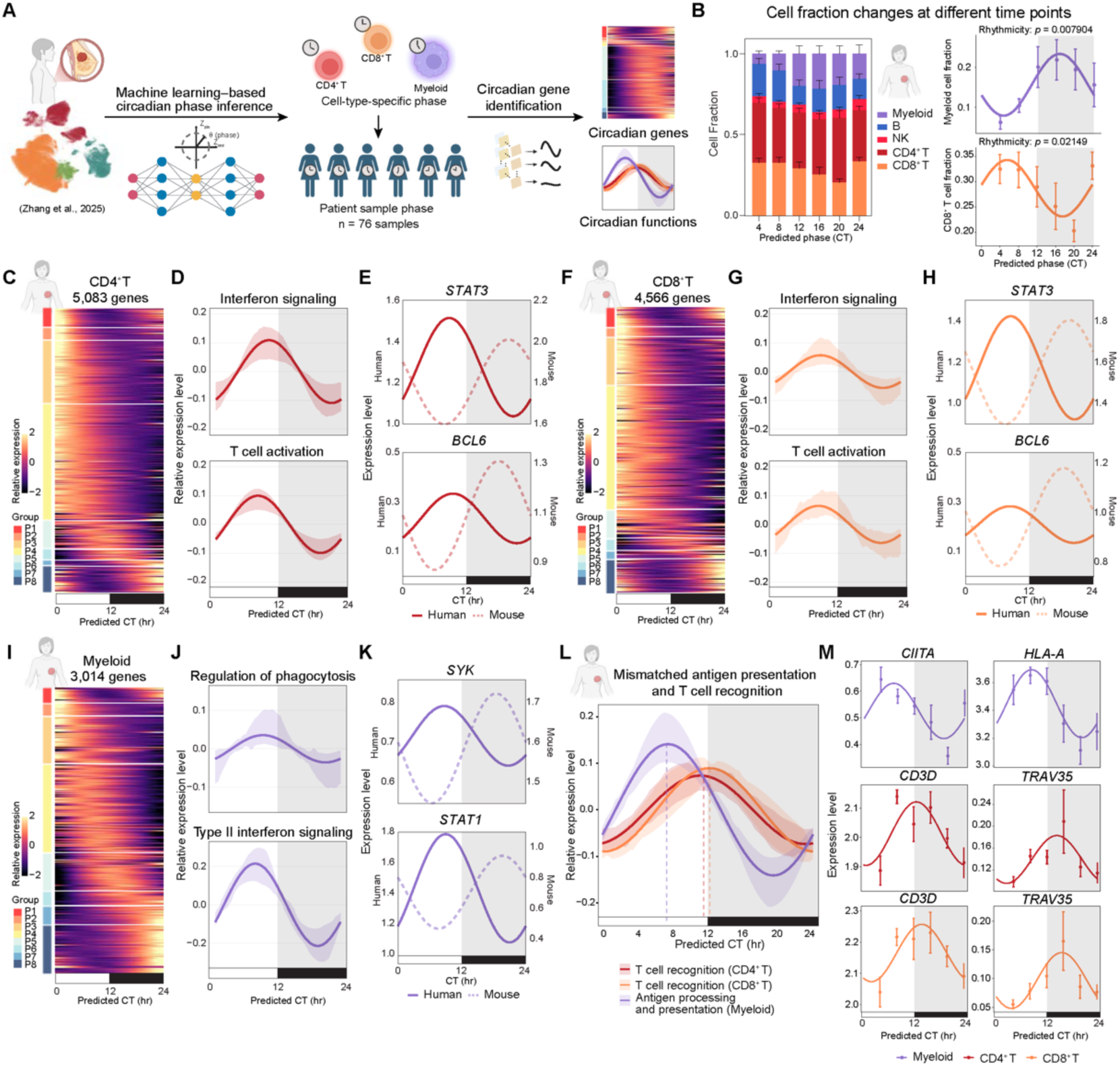
Circadian regulation in human TNBC. (A) Workflow illustrating circadian phase inference and rhythmic gene identification in human TNBC single-cell RNA-seq data^23^ using CYCLOPS2.0 and DiscoRhythm. (B) Left: Stacked bar plots showing relative proportions of immune cell populations across predicted circadian phases within the human TNBC tumor microenvironment (TME). Right: Line plots showing circadian fitting curves of myeloid (top) and CD8⁺ T cell (bottom) fractions across a circadian day. Points indicate mean values at each time point (± SEM). Gray shading denotes the dark phase (CT12–CT24/CT0). *p* values indicate rhythmicity assessed by cosinor regression. *n* = 10-13 patient samples per time point. (C) Heatmap showing rhythmic expression patterns of circadian genes in CD4⁺ T cells across inferred circadian phases. The color bar reflects the expression level from high (bright yellow) to low (deep purple). (D) Circadian fitting curves showing the rhythmic dynamics of representative functional pathways in CD4⁺ T cells, including interferon signaling and T cell activation. Bold lines represent mean fitted curves across genes within each pathway, shaded areas indicate the interquartile range (25th–75th percentile), and the gray area indicates the dark phase (CT12–CT24/CT0). (E) Circadian fitting curves of representative genes (*STAT3* and *BCL6*) in CD4⁺ T cells. Solid lines indicate human TNBC data, and dashed lines indicate the corresponding mouse tumor data. The gray area indicates the dark phase (CT12–CT24/CT0). (F) Heatmap showing rhythmic expression patterns of circadian genes in CD8⁺ T cells across inferred circadian time points. The color bar reflects the expression level from high (bright yellow) to low (deep purple). (G) Circadian fitting curves showing the rhythmic dynamics of representative functional pathways in CD8⁺ T cells, including interferon signaling and T cell activation. Bold lines represent mean fitted curves across genes within each pathway, shaded areas indicate the interquartile range (25th–75th percentile), and the gray area indicates the dark phase (CT12–CT24/CT0). (H) Circadian fitting curves of representative genes (*STAT3* and *BCL6*) in CD8⁺ T cells. Solid lines indicate human TNBC data, and dashed lines indicate the corresponding mouse tumor data. The gray area indicates the dark phase (CT12–CT24/CT0). (I) Heatmap showing rhythmic expression patterns of circadian genes in myeloid cells across inferred circadian time points. The color bar reflects the expression level from high (bright yellow) to low (deep purple). (J) Circadian fitting curves showing the rhythmic dynamics of representative functional pathways in myeloid cells, including regulation of phagocytosis and type II interferon signaling. Bold lines represent mean fitted curves across genes within each pathway, shaded areas indicate the interquartile range (25th–75th percentile), and the gray area indicates the dark phase (CT12–CT24/CT0). (K) Circadian fitting curves of representative genes (*SYK* and *STAT1*) in myeloid cells. Solid lines indicate human TNBC data, and dashed lines indicate the corresponding mouse tumor data. The gray area indicates the dark phase (CT12–CT24/CT0). (L) Circadian fitting curves illustrating the temporal mismatch between antigen processing and presentation in myeloid cells and T cell recognition programs in CD4⁺ and CD8⁺ T cells. Bold lines represent mean fitted curves across genes within each pathway, shaded areas indicate the interquartile range (25th–75th percentile). Vertical dashed lines indicate the peak timing of each functional program. The gray area indicates the dark phase (CT12–CT24/CT0). (M) Circadian expression profiles of representative genes involved in antigen presentation (*CIITA*, *HLA-A*) in myeloid cells and T cell recognition (*CD3D*, *TRAV35*) in CD4⁺ and CD8⁺ T cells. Points represent mean expression values at each inferred circadian time point, with error bars indicating SEM. The gray area indicates the dark phase (CT12–CT24/CT0).

With circadian phase established, we further identified cell-type-specific circadian genes and pathways in CD4⁺ T cells, CD8⁺ T cells, and myeloid populations, enabling downstream analyses of immune circadian organization in human TNBC. Similar to our findings in the mouse model, circadian phases of CD4⁺ T cells, CD8⁺ T cells, and myeloid cells were highly concordant in human TNBC, indicating preserved immune circadian coordination (**Figure S8D**). Core clock genes such as *ARNTL*, *NR1D2*, *PER2*, and *DBP* showed similar relative phase ordering across immune cell types, in agreement with prior observations in diurnal primate models,^56^ supporting the validity of circadian phase inference in humans (**Figures S8E and S8F**).

Comparative circadian analysis identified 242, 85, and 179 genes in CD4⁺ T cells, CD8⁺ T cells, and myeloid cells, respectively, that exhibited conserved but phase-shifted circadian regulation in human TNBC relative to the mouse tumor model (**Figure S8G**). Functional enrichment analysis revealed that these genes were associated with T cell activation, interferon signaling, and T cell receptor signaling in CD4⁺ and CD8⁺ T cells (e.g., *BCL6*, *RUNX3*, *STAT3*), and with phagocytosis, interferon signaling, and antigen processing and presentation in myeloid cells (e.g., *SYK*, *STAT1*, *TAP1*, *TAPBP*) (**Figure S8G**). In human TNBC, these immune activation–related pathways predominantly peak during the inferred daytime (**Figures 7C–7K and S8H–S8J**). These results define a daytime-associated (active phase) immune-active state in human TNBC, which may help explain clinical observations that antitumor immune responses and immunotherapy efficacy are enhanced during daytime phases.^13,15,57–59^ However, similar to our observations in mouse tumors, we found a circadian mismatch in human TNBC between the peak timing of antigen processing and presentation in myeloid cells and T cell recognition programs in CD4⁺ and CD8⁺ T cells (**Figures 7L and 7M and S8K**). This temporal misalignment indicates an uncoupling between antigen availability and T cell responsiveness within the TME, which potentially impairs optimal antitumor immune coordination and facilitates immune evasion.

Together, these findings uncover a conserved circadian crosstalk among TME compartments in both mouse and human tumors, demonstrating that temporal coordination and misalignment critically influence immune surveillance within tumors. These results position circadian regulation as a fundamental axis of tumor–immune dynamics and suggest opportunities to leverage biological timing for improved cancer immunotherapy.

## DISCUSSION

Circadian rhythms regulate diverse physiological programs within immune and cancer cells.^60,61^ Yet, how these intrinsic oscillations are organized across cellular populations within the tumor microenvironment, and whether intercellular rhythmic regulation shapes the functional architecture of the TME and influences tumor development and treatment, remain largely undefined. Here, using time-resolved single-nucleus multiomic profiling, we delineate the circadian regulatory architecture of the TNBC tumor microenvironment, encompassing proliferation, immune responses, and metabolic programs across diverse cellular populations. We identify three principal axes of circadian misalignment across the tumor microenvironment. First, temporal decoupling between tumor proliferation and immune activation restricts effective immune engagement. Second, discordance between antigen presentation and T cell recognition limits immune surveillance efficiency, while overlapping activation and exhaustion-associated programs further constrain T cell effector function. Third, asynchronous oscillations of PD-1 and PD-L1 sustain checkpoint-mediated suppression across circadian phases. Together, these findings establish immune escape as a temporally structured and dynamically misaligned process, providing a conceptual framework for considering temporal organization in therapeutic strategy.

Recent single-cell profiling studies have revealed extensive cellular heterogeneity within the breast cancer tumor microenvironment, uncovering diverse cellular states and intercellular interactions that drive tumor progression and shape therapeutic response.^22–30,62^ However, these analyses are largely based on single-timepoint measurements that portray temporally dynamic and circadian-regulated programs as static cellular states. Our time-resolved single-cell multiomic analyses extended this framework and introduced a temporal regulatory dimension to the TME landscape. We identified both shared and cell-type-specific circadian genes that coalesced into higher-order functional modules across the TME. Leveraging chromatin accessibility profiles, we further inferred transcription factor activity to delineate the regulatory logic underlying these rhythmic processes. These analyses revealed that fundamental programs governing cellular proliferation, immune function, tissue organization, RNA processing and translation, and metabolic pathways were embedded within circadian control. Moreover, drug-associated enrichment analyses indicated that several circadian modules harbored targets of therapeutic agents, including paclitaxel. Notably, many of these targets also oscillated in immune populations, suggesting that therapeutic effects may vary across circadian phases. Together, these findings underscore that incorporating temporal resolution into single-cell analyses establishes a more comprehensive framework for understanding circadian dynamics within the TME and suggests that temporal organization may shape both tumor development and therapeutic response.

Breast cancers comprise heterogeneous epithelial states associated with distinct functional programs,^28,63,64^ yet whether these states undergo circadian remodeling within tumors has remained unclear. We identified two cancer epithelial populations, a proliferative state and an inflammatory state, that rhythmically transitioned across the circadian cycle *in vivo*. Notably, these circadian epithelial states were conserved in human TNBC. Our findings indicated that subtype composition and intrinsic rhythmicity jointly shaped cancer epithelial circadian programs, with proliferative activity peaking in the early daytime and antigen-presentation programs reaching maximal levels at late night. Consistent with our findings, prior studies have reported time-of-day–dependent increases in tumor cell dissemination and expansion in murine models.^10,65^ Similarly, immune components within the TNBC TME exhibited robust circadian organization shaped by both oscillations in subtype composition and intrinsic rhythmic regulation within each subtype. CD4⁺ and CD8⁺ T cells displayed peak activation and effector programs during the night, corresponding to the murine active phase. The macrophage/monocyte compartment exhibited a similar temporal pattern, with phagocytosis, antigen presentation, and interferon-response programs peaking at night. Retrospective clinical studies in metastatic non–small cell lung cancer and melanoma have reported time-of-day differences in the efficacy of immune checkpoint inhibitors, with morning administration associated with improved survival outcomes.^13,15,57,58,66–72^ Our findings provided a single-cell and multi-compartment framework that offers mechanistic insight into these time-dependent differences in therapeutic response. Consistent with our findings, a mechanistic study in a mouse melanoma model demonstrated that rhythmic tumor infiltration and cytotoxic T cell activation peaked during the active phase, enhancing the efficacy of time-restricted anti–PD-1 therapy.^53^ Together, these observations indicate that both tumor-intrinsic and immune effector programs are temporally organized, positioning circadian phase as an important determinant of tumor–immune dynamics.

Whether such rhythmic programs are temporally coordinated across cellular compartments within the tumor microenvironment has remained unclear. Our phase-resolved single-cell analyses revealed that while functionally related immune populations maintained strong phase concordance, cancer epithelial cells exhibited broad phase divergence relative to stromal and immune compartments, defining three principal axes of circadian misalignment in TNBC. First, proliferative programs in cancer epithelial cells peaked during the early-daytime, whereas effector and interferon-response pathways in T cells and macrophages reached maximal activation at night. This phase separation temporally decoupled tumor cell division from peak immune effector function, reducing the temporal efficiency of antitumor responses.

Second, antigen-presentation programs in cancer epithelial and myeloid cells peaked at night, whereas T cell receptor signaling pathways peaked during the day, indicating a temporal separation between antigen availability and immune recognition. Moreover, activation-associated programs in T cells were phase-concordant with inhibitory receptors and exhaustion markers, resulting in the concurrent engagement of stimulatory and suppressive pathways. Such activation–inhibition coupling may constrain effective immune responses and favor differentiation toward dysfunctional states, consistent with prior studies demonstrating that persistent co-engagement of activating and inhibitory signaling promotes T cell exhaustion.^73,74^

Third, PD-1 and PD-L1 rhythms were not synchronized across cellular compartments. The PD-1/PD-L1 checkpoint axis is a central regulator of T cell activation and tumor immune evasion and a major therapeutic target in cancer.^19–21,50–52,75–77^ We found that PD-1 protein levels in T cells peaked during the early night, consistent with prior melanoma studies showing circadian regulation of PD-1 expression.^53^ In contrast, PD-L1 expression in cancer epithelial and macrophage populations peaked during the early daytime, establishing a phase mismatch across compartments. This temporal misalignment suggests that inhibitory signaling may persist even as T cells transition toward a more activated state. Collectively, these three axes of circadian misalignment reposition tumor–immune interactions as a circadian-dependent system in which temporal decoupling across cellular compartments constrains effective antitumor immunity.

Human studies are often limited by the absence of precise sampling times, precluding systematic assessment of circadian immune regulation. In this study, we reconstructed temporal immune dynamics and identified rhythmic gene expression across CD4⁺ T, CD8⁺ T, and myeloid populations by applying CYCLOPS 2.0 to infer circadian phase in human TNBC single-nucleus RNA-seq datasets.^55,78^ Similar to the murine TME, immune activation and macrophage phagocytosis in human TNBC peaked during the active phase. Moreover, genes involved in antigen processing in myeloid cells and T cell recognition programs in CD4⁺ and CD8⁺ T cells exhibited circadian phase misalignment, revealing a similar temporal architecture across species despite reversed light–dark cycles.

Recent retrospective clinical studies in melanoma and non–small cell lung cancer have reported that the efficacy of immune checkpoint blockade varies according to the circadian time of administration, with morning dosing associated with improved survival outcomes.^13–15,57,79^ Notably, given the prolonged half-life of therapeutic antibodies,^80,81^ these findings raise the possibility that the circadian phase at the time of initial tumor–antibody encounter may critically shape early immune remodeling within the TME. Although such timing effects have not yet been systematically examined in TNBC, our data provide a mechanistic framework for considering circadian organization as a variable in TNBC immunotherapy. Together, our findings establish temporal organization as an integral dimension of tumor–immune biology and suggest that incorporating circadian phase into therapeutic design may represent a previously underappreciated avenue for optimizing immunotherapy in breast cancer.

### Limitations of the study

Effective tumor–immune communication likely depends not only on temporal coordination but also on spatial organization within the tumor microenvironment. Beyond synchronized timing, rhythmic programs are shaped by how distinct cellular populations are positioned and interact across the tissue landscape. Incorporating spatial transcriptomics and multiplex imaging with temporal profiling would provide a more integrated understanding of how local architecture, cell–cell signaling, and microenvironmental gradients shape circadian coordination or disruption within the TME. In addition, the circadian rhythms observed in animal models reflect an idealized system under controlled conditions. In contrast, human circadian regulation is shaped by diverse physiological, environmental, and behavioral factors, leading to considerable inter-individual variability. For clinical translation, it will be critical to develop computational methods that integrate molecular, physiological, and lifestyle cues to infer each patient’s circadian rhythm. Such approaches may enable personalized time-based immunotherapy with improved efficacy.

## RESOURCE AVAILABILITY

### Lead contact

Further information and requests for resources and reagents should be directed to and will be fulfilled by the lead contact Myles Brown.

### Materials availability

Materials and reagents can be requested from the lead contact upon reasonable request. This study did not generate new unique reagents.

### Data and code availability

All sequencing data generated in this study have been deposited in GEO and will be publicly released as of the date of publication

All original code has been deposited on GitHub and will be publicly released as of the date of publication.

Any additional information required to reanalyze the data reported in this paper is available from the lead contact upon request.

## ACKNOWLEDGMENTS

This study was supported by the Claudia Adams Barr Program at Dana-Farber Cancer Institute and the Ludwig Center at Harvard. We thank Shuqiang Li, PhD and Wesley S. Lu from Dana-Farber Cancer Institute Translational Immunogenomics Laboratory for their technical assistance with the snMultiome. We thank the Dana-Farber Cancer Institute Medical Oncology, DanaFarber/Harvard Cancer Center Rodent Histopathology Core facility, Dana-Farber Cancer Institute Animal Resource Facilities, Translational Immunogenomics Laboratory and Neurobiology Imaging Facility at Havard Medical School for outstanding services.

## AUTHOR CONTRIBUTIONS

M.B., M.K., C.L., and Z.L. conceived the project and designed the experiments. C.L. and Z.L. carried out the experiments. C.L., Z.L., and X.X. performed the bioinformatic analysis of the snMultiomic data. Z.X., C.L., and Z.L. performed human sample phase prediction analyses. Q.Z., T.Z., B.X., and G.W. assisted with mouse tissue collection. C.L., Z.L., and X.X. wrote the manuscript. All authors reviewed and edited the manuscript. M.B. and M.K. supervised the work.

## DECLARATION OF INTERESTS

M.B. receives sponsored research support from Novartis and serves on the scientific advisory board of GV20 Therapeutics and has equity in the company. R.J. received research funding from Lilly, Pfizer and Novartis and serves on an advisory board for GE Healthcare and Carrick Therapeutics. The other authors declare no competing interests.

## DECLARATION OF GENERATIVE AI AND AI-ASSISTED TECHNOLOGIES

During the preparation of this manuscript, the authors used OpenAI’s GPT-5.2 model for language editing and proofreading to improve clarity and readability. The authors reviewed and revised all output as needed and took full responsibility for the content of the publication.

## STAR METHODS

### KEY RESOURCES TABLE

| REAGENT or RESOURCE | SOURCE | IDENTIFIER |
| --- | --- | --- |
| Experimental models: Organisms/strains |  |  |
| Mouse: BALB/cJ | The Jackson Laboratory | Strain #:000651; RRID:IMSR_JAX:000651 |
| Antibodies |  |  |
| Anti-mouse Ki-67 antibody | R&D | Cat# AF7649; RRID: AB_2687500 |
| Anti-mouse beta-2 Microglobulin antibody | Thermo Fisher Scientific | Clone 4H5L6; RRID:AB_2532441 |
| Anti-mouse pan Cytokeratin antibody | Abcam | Clone C11; RRID:AB_306047 |
| Anti-mouse PD-1 | Cell Signaling Technology | Clone D7D5W;RRID:AB_2800041 |
| Anti-mouse CD3 | Cell Signaling Technology | Clone 17A2; RRID:AB_2800082 |
| Anti-mouse PD-L1 | Cell Signaling Technology | Clone F3Z1T; RRID: AB_3730179 |
| Anti-mouse F4/80 | Cell Signaling Technology | Clone BM8.1; RRID: AB_2938669 |
| Chemicals, peptides, and recombinant proteins |  |  |
| Protector RNase Inhibitor | Roche Diagnostics | Cat# 3335402001 |
| DL-Dithiothreitol solution | Sigma Aldrich | Cat# 646563 |
| Phosphate-Buffered Saline (PBS) | Fisher Scientific | SH3025601 |
| IGEPAL® CA-630 for molecular biology | Sigma-Aldrich | Cat# I8896 |
| Digitonin (5%) | Life Technologies | Cat # BN2006 |
| Heparin sodium | Selleck | Cat# S1346 |
| CD45 (TIL) MicroBeads, mouse | Miltenyi Biotec | Cat # 130-110-618 |
| Antigen Retrieval Buffer | Abcam | Cat# ab93684 |
| Critical commercial assays |  |  |
| Chromium Next GEM Single Cell Multiome ATAC + Gene Expression Kit | 10x Genomics | Cat# 1000283 |
| Library Construction Kit | 10x Genomics | Cat# 1000190 |
| Chromium Nuclei Isolation with RNase Inhibitor Kit | 10x Genomics | Cat# 1000494 |
| Deposited data |  |  |
| Processed snMultiome data in mouse model | This study | https://compbio.mit.edu/Circadian_BRCA/ |
| Processed scRNA-seq data from TNBC patients | Zhang et al.<sup>23</sup> | GEO: GSE266919 |
| Processed scRNA-seq data from BRCA patients | Wu et al.<sup>28</sup> | GEO: GSE176078 |
| Processed snRNA-seq data from BRCA patients | Klughammer et al.<sup>49</sup> | dbGaP: phs002371 (https://humantumoratlas.org) |
| Software and algorithms |  |  |
| cellranger-arc | 10x Genomics | https://support.10xgenomics.com/single-cell-atac/software |
| CellBender (v.0.3.2) | Stephen et al.<sup>82</sup> | https://github.com/broadinstitute/CellBender |
| Scanpy (v.1.9.3) | Wolf et al.<sup>83</sup> | https://scanpy.readthedocs.io/en/stable/ |
| scrublet (v.0.2.1) | Wolock et al.<sup>84</sup> | https://github.com/swolock/scrublet |
| Muon (v.0.1.3) | Bredikhin et al.<sup>85</sup> | https://muon.readthedocs.io/en/latest/index.html |
| Seurat (v.4.4.0) | Hao et al.<sup>86</sup> | https://satijalab.org/seurat/ |
| ArchR (v.1.0.2) | Granja et al.<sup>87</sup> | https://www.archrproject.com/ |
| MACS2 (v.2.2.6) | Zhang et al.<sup>88</sup> | https://pypi.org/project/MACS2/ |
| GREAT (v.1.26.0) | McLean et al.<sup>89</sup> | http://great.stanford.edu/ |
| ComplexHeatmap (v.2.14.0) | Gu et al.<sup>90</sup> | https://jokergoo.github.io/ComplexHeatmap-reference/book/ |
| scCustomize (v.2.1.1) | Marsh et al.<sup>91</sup> | https://github.com/samuel-marsh/scCustomize/ |
| ggpubr (v.0.6.0) | Kassambara et al. | https://github.com/kassambara/ggpubr |
| IGV Genome Browser (v.2.15.4) | Robinson et al.<sup>92</sup> | https://igv.org/ |
| pheatmap (v.1.0.12) | Kolde et al.<sup>93</sup> | https://github.com/raivokolde/pheatmap |
| enrichR (v.3.2) | Kuleshov et al.<sup>94</sup> | https://maayanlab.cloud/Enrichr/ |
| BEDTools (v.2.30.0) | Quinlan et al.<sup>95</sup> | https://bedtools.readthedocs.io/en/latest/ |
| ggplot2 (v.3.4.0) | Randle Aaron et al.<sup>96</sup> | https://ggplot2.tidyverse.org/ |
| DiscoRhythm | Carlucci et al.<sup>31</sup> | https://bioconductor.org/packages/release/bioc/html/DiscoRhythm.html |
| Metascape | Zhou et al.<sup>97</sup> | https://metascape.org/gp/index.html#/main/step1 |
| Cytoscape (v.3.10.4) | Shannon et al.<sup>98</sup> | https://cytoscape.org/ |
| QuPath (v.0.4.0) | Bankhead et al.<sup>99</sup> | https://qupath.readthedocs.io/en/0.4/index.html |
| CYCLOPS 2.0 | Li et al.<sup>55</sup> | https://researchdiscovery.drexel.edu/esploro/outputs/doctoral/CYCLOPS-20-improving-and-applying-circadian/991022064341804721 |
| chromVAR | Schep et al.<sup>100</sup> | http://www.github.com/GreenleafLab/chromVAR |
| inferCNV (v.1.12.0) | Broad institute | https://github.com/broadinstitute/inferCNV |

### EXPERIMENTAL MODEL AND STUDY PARTICIPANT DETAILS

#### Animals

Female BALB/cJ mice were purchased from The Jackson Laboratory and housed under a 12 h:12 h light:dark (LD) cycle with ad libitum access to food and water. For the breast cancer model, 5 × 10⁵ 4T1 cells in 100 μL of PBS were orthotopically injected into the mammary fat pad of 10-week-old female BALB/cJ mice under isoflurane anesthesia. To assess circadian dynamics, tumor-bearing mice were maintained under LD conditions for 10 days to allow tumor establishment, followed by a 24-hour period of constant darkness (DD) for circadian adaptation. Tumors were harvested under dark conditions at four circadian time (CT) points: CT4, CT10, CT16, and CT22. All animal experiments were conducted in a specific pathogen–free (SPF), rodent-only barrier facility accredited by AAALAC at Dana-Farber Cancer Institute (Boston, MA), in strict accordance with protocol 09-044 approved by the Dana-Farber Cancer Institute Institutional Animal Care and Use Committee (IACUC).

### METHOD DETAILS

#### Tumor collection and cell isolation

Tumors were dissected, weighed, and mechanically dissociated, followed by enzymatic digestion with Collagenase/Hyaluronidase (StemCell Technologies, #07912) at 37°C with gentle shaking. Digested tissues were passed through a 70 μm cell strainer, washed, and treated with red blood cell lysis buffer (Thermo Fisher Scientific, #BDB555899). After neutralization and additional washes, single-cell suspensions were incubated with CD45 MicroBeads (Miltenyi Biotec, #130-052-301) according to the manufacturer’s protocol. MACS LS columns were pre-equilibrated with MACS buffer (0.5% BSA + 2 mM EDTA in PBS) and mounted on a MidiMACS™ separator. The labeled cell suspension was loaded onto the column, and the CD45⁻ fraction was collected in the flow-through after two washes with MACS buffer. The column was then removed from the magnetic stand, and CD45⁺ cells were eluted using 3 mL of MACS buffer with a plunger.

#### Single nucleus multiome library construction and sequencing

To isolate nuclei for multiomic-seq, we followed the manufacturer’s protocol with modifications (10x Genomics, CG000365 • Rev C). Cryopreserved cells were rapidly thawed in a 37°C water bath until a small ice crystal remained, then transferred to a 50 mL conical tube. The cryovial was rinsed with pre-warmed RPMI supplemented with 10% FBS and added dropwise to the tube with gentle mixing. Cells were gradually diluted by five sequential 1:1 additions of media, with 1-minute intervals between steps. After centrifugation (300 rcf, 5 min), the cell pellet was resuspended in 0.04% BSA in PBS, washed, and filtered through a 40 μm Flowmi strainer. For nuclei isolation, cells were then centrifuged (300 rcf, 5 min, 4°C), lysed in 100 μL of chilled lysis buffer (10x Genomics) by gentle pipetting, and incubated on ice for 5 minutes. Lysis was stopped by washing with a chilled wash buffer three times. Nuclei were resuspended in chilled Diluted Nuclei Buffer (10x Genomics) at a concentration of 5,000 nuclei/μL and immediately used for multiome library preparation. RNase inhibitor (Roche Diagnostics, #3335402001) was included in all buffers.

We performed simultaneous profiling of single-nucleus transcriptomes and chromatin accessibility using Chromium Next GEM Single Cell Multiome ATAC + Gene Expression Reagent Bundle (10x Genomics, PN-1000283). For each sample, about 10,000 nuclei were bulk-transposed and loaded onto the 10x Chromium X system for GEM (Gel Bead-in-Emulsion) generation and barcoding. Following pre-amplification PCR, the material was split to separately construct single-nucleus ATAC-seq (snATACseq) and gene expression (GEX) libraries. Final snATAC and snGEX libraries were independently pooled, and sequenced on an Illumina NovaSeq 6000 S2 (snATACseq) and S4 flow cell (GEX), using the manufacturer’s recommended read lengths and sequencing depth for each library type.

#### Hematoxylin and Eosin (H&E) staining

For histological analysis, tumor were fixed with 10% formalin for 24 hours. Samples were then paraffin-embedded and sectioned at a thickness of 5 μm. Tissue sections were deparaffinized in xylene and rehydrated through a descending ethanol gradient (100%, 100%, 95%, 90%, 80%, 70%, 50%), followed by a rinse in distilled water. Sections were stained with hematoxylin for 5 minutes, washed under running water, differentiated using 1% acid alcohol, and rinsed again until nuclei staining appeared optimal. Subsequently, sections were counterstained with eosin, dehydrated through an ascending ethanol series (50%, 70%, 80%, 90%, 95%, 100%, 100%), and immersion in xylene. Slides were then coverslipped using neutral balsam for long-term storage.

#### Immunofluorescence staining

For immunofluorescence staining of paraffin-embedded tumor sections, sections were deparaffinized, rehydrated, and subjected to antigen retrieval in Tris-EDTA Buffer, pH 9.0 (Abcam, ab93684). After cooling, sections were blocked with 10% donkey serum in PBS or the blocking buffer from the Mouse on Mouse (M.O.M.) Immunodetection Kit (Vector Laboratories, BMK-2202) for 60 minutes at room temperature. Primary antibodies were then applied at appropriate dilutions and incubated overnight at 4°C. Following PBS washes, sections were incubated with secondary antibodies diluted in 10% donkey serum for 1 hour at room temperature, protected from light. Sections were washed in PBS, counterstained with DAPI for 5 minutes at room temperature, washed again in PBS, and mounted using ProLong™ Gold Antifade Mountant (Invitrogen, P36930).

For immunofluorescence staining of frozen sections, tumor tissues were embedded in OCT compound and cryosectioned. Sections were air-dried, fixed in 4% paraformaldehyde for 10 minutes at room temperature, and washed in PBS. Sections were then blocked for 60 minutes at room temperature with either 10% donkey serum in PBS or the blocking buffer from the Mouse on Mouse (M.O.M.) Immunodetection Kit (Vector Laboratories, BMK-2202). Primary antibodies were applied at appropriate dilutions in 1% donkey serum in PBS and incubated overnight at 4°C. Following PBS washes, sections were incubated with fluorescently labeled secondary antibodies diluted in 1% donkey serum in PBS for 1 hour at room temperature, protected from light. Sections were washed in PBS, counterstained with DAPI for 5 minutes at room temperature, washed again in PBS, and mounted using ProLong™ Gold Antifade Mountant (Invitrogen, P36930).

Images were acquired using a Zeiss LSM 980 confocal microscope and analyzed with QuPath (version 0.4.0). Circadian rhythmicity of protein expression was assessed by 24-h cosinor regression implemented in R, modeling values as a linear combination of sine and cosine terms sin(2πt/24) and cos(2πt/24). Statistical significance was evaluated by an F-test comparing the full model to a constant model, and fitted curves were generated from the estimated parameters.

### QUANTIFICATION AND STATISTICAL ANALYSIS

#### Preprocessing and quality control of snMultiome data

Raw data generated using the 10x Genomics Multiome platform were processed with the *cellranger-arc count* toolkit (v.2.0.2) following the manufacturer’s instructions and using the GRCm38 (mm10) mouse reference genome. This pipeline produced fragment files containing transposase cut sites from snATAC-seq data, as well as filtered and aligned gene expression matrices from snRNA-seq data. The resulting raw gene expression count matrices generated by Cell Ranger were subsequently processed to remove ambient RNA contamination using the CellBender *remove-background* function (v.0.2.2) with default parameters.^82^ Downstream analyses of snATAC-seq and snRNA-seq data, including dimensionality reduction, clustering, peak calling and annotation, identification of cell-type-specific genes, and TF analyses were performed using Scanpy (v.1.9.3), Muon (v.0.1.3), and ArchR (v.1.0.2).^83,85,87^

ArchR (v1.0.2) was used to process fragment files generated from snATAC-seq data, producing Arrow files and ArchR projects for downstream analyses. Quality control of snATAC-seq nuclei was performed in ArchR, including doublet detection and removal using the *filterDoublets* function. Nuclei with transcription start site (TSS) enrichment scores greater than 2 and more than 1,000 fragments were retained. Fragment size distributions across all snATAC-seq libraries were assessed using the *plotFragmentSizes* function. In parallel, Muon (v.0.1.3) was used to process fragment files and generate AnnData (*.h5ad*) objects.^85^

For clustering analysis of snATAC-seq data, we performed iterative clustering based on snATAC profiles. In the initial round, dimensionality reduction and clustering were conducted in ArchR using the *addIterativeLSI* and *addClusters* functions with default parameters.^87^ After preliminary cell-type annotation, clustering was further refined using a union peak set defined across cell types in Muon (v.0.1.3).^85^ Accessible fragments overlapping each peak were quantified for each nucleus using *muon.atac.tl.locate_fragments* and *muon.atac.tl.count_fragments_features*. The resulting tile matrix was normalized using term frequency–inverse document frequency (TF–IDF), followed by latent semantic indexing (LSI) on the scaled matrix to derive latent components. Consistent with established practices, the first LSI component, typically correlated with technical factors such as sequencing depth and fragment counts, was excluded from downstream analyses. Two-dimensional visualization of snATAC-seq nuclei was performed using UMAP as implemented in Scanpy (v.1.9.3) via the *scanpy.tl.umap* function, using the LSI embeddings.^83^

Cellbender-processed snRNA-seq data were processed and integrated using Scanpy (v.1.9.3),^83^ including quality control, normalization, dimensionality reduction, and clustering. Doublets were identified and removed using Scrublet (v.0.2.1) as implemented in Scanpy.^84^ Nuclei with fewer than 200 detected genes, greater than 10% mitochondrial reads were excluded. The remaining nuclei were processed following the standard Scanpy workflow, including total count normalization, log transformation, and principal component analysis (PCA) using *scanpy.pp.pca*. Neighbor graph and UMAP embedding were then computed, and cell clusters were identified using the Leiden algorithm *scanpy.tl.leiden*. To achieve high-confidence cell type clustering and annotation, a second round of quality control and doublet removal was performed within each cell type. Cells were subset by cell type and reprocessed through normalization, dimensionality reduction, and UMAP embedding to define the cell subtypes. Clusters exhibiting abnormal read counts, atypical numbers of detected genes, or ambiguous marker gene expression were excluded from downstream analyses.

Only high-quality nuclei passing quality control for both snRNA-seq and snATAC-seq modalities were retained for downstream analyses.

#### Annotation of the cell types and subtypes

Multiple complementary approaches were used to annotate cell types and subtypes in the snRNA-seq data. Cell types and subtypes were initially identified based on the expression of established canonical marker genes specific to each cell type, supported by gene expression patterns in snRNA-seq data and corresponding gene activity scores inferred from snATAC-seq profiles.^24,27,28,62,101–112^ Marker genes including *Cd4*, *Lef1*, and *Cd3d* (CD4⁺ T cells), *Cd8a* and *Cracr2a* (CD8⁺ T cells), *Cd8b*, *Arhgap9*, and *Lcn4* (NKT cells), *Ncr1* and *Nkg7* (NK cells), *Cd19* and *Ms4a1* (B cells), *Lyz2*, *Csf1r*, and *C1qa* (Macro/Mono), *S100a9*, *Retnlg*, and *Csf3r* (neutrophils), *Flt3* and *Clec9a* (DCs), *Col1a1*, *Col3a1*, and *Col5a2* (CAFs), *Robo4*, *Cdh5*, and *Ptprb* (endothelial cells), *Rgs5* and *Abcc9* (PVLs), *Krt5* and *Elf5* (normal epithelial cells), *Hmga2*, *Sox9*, *Epcam*, *Krt18*, and *Krt8* (cancer epithelial cells), and *Jchain*, *Mzb1*, and *Xbp1* (plasmablasts).

Beyond major cell types, cell subtypes were subsequently identified based on established subtype-specific marker genes reported in prior studies.^27,28,113–115^ To further refine cell subtype annotations, we applied gene set-based scoring approaches. Functional gene module scores were computed using the *AddModuleScore* function implemented in Seurat, enabling the identification of transcriptional programs associated with specific biological processes or cell states. In addition, to distinguish proliferative and inflammatory cancer cell subtypes, cell cycle phase scores were calculated using Seurat’s *CellCycleScoring* function, allowing classification of nuclei according to cell cycle status. Gene signatures used for scoring were derived from previously published studies and listed in Table S4.^23,116^

#### Identification of the cell-type-specific genes and TFs

Cell-type-specific genes for the 14 identified cell types were defined based on both gene expression profiles from snRNA-seq data and gene activity scores derived from snATAC-seq data. Differential gene expression analysis was performed using the *sc.tl.rank_genes_groups* function in Scanpy,^83^ while differential analysis of gene activity scores was conducted using the *getMarkerFeatures* function in ArchR.^87^ Genes with an FDR < 0.05 and log_2_ (fold change) > 1 in both snRNA-seq and snATAC-seq analyses were considered statistically significant cell-type-specific genes. To identify cell-type-specific TFs, we first detected cell-type-specific cCREs by applying the Wilcoxon rank-sum test to the peak matrix using the *getMarkerFeatures* function in ArchR.^87^ cCREs with FDR < 0.1 and log_2_ (fold change) > 0.5 were retained and subsequently annotated with TF motifs using the *peakAnnoEnrichment* function. The resulting motif enrichments were visualized using the *plotEnrichHeatmap* function.

#### Circadian genes and gene modules

To identify circadian genes in each cell type based on snRNA-seq data, we performed circadian rhythmicity analysis using the DiscoRhythm R package (v.1.10.1).^31^ Genes exhibiting significant 24-hour periodic expression patterns were classified as circadian. The normalized expression matrix was aggregated into a pseudo-bulk matrix across replicates and experimental time points (CT4, CT10, CT16, and CT22) for each cell type. For each cell type, low expression genes were filtered by total expression values greater than 0.2 across all pseudo-bulked samples. The resulting data matrix was analyzed using the built-in diagnostic functions *discoInterCorOutliers* and *discoPCAoutliers* to detect and remove outliers, with all parameters set to default. Gene selection was performed using the *discoRepAnalysis* function, which applies ANOVA statistical testing across biological replicates. Genes with p < 0.05 and F-statistics > 1 were retained for subsequent analyses. The primary circadian period of interest (24 hours) was evaluated using the *discoPeriodDetection* function, followed by fitting a Cosinor model to characterize rhythmic oscillations. Finally, oscillatory genes were identified using the *discoODAs* function with the Cosinor (CS) method. Circadian genes identified in each cell type are provided in Table S1.

The identified circadian genes were further analyzed to identify circadian gene modules. The normalized cell-by-gene expression matrix was subset to include only circadian genes and then transposed to generate a gene-by-cell matrix. Dimensionality reduction was performed using principal component analysis (PCA) implemented in Scanpy.^83^ For visualization, the PCA-transformed gene expression profiles were projected onto a UMAP embedding. Circadian genes were subsequently clustered using the Leiden *sc.tl.leiden* algorithm, resulting in 20 circadian gene modules. Circadian genes within each module are listed in Table S2.

#### Functional and drug target enrichment of circadian gene modules

Circadian genes within each module were analyzed for functional pathway enrichment using Metascape with default parameters, incorporating Gene Ontology (GO), KEGG, Reactome, and WikiPathways databases.^97^ Enriched terms with adjusted *p* values < 0.05 were considered significant. Drug target enrichment was performed using enrichR (v.3.2),^94^ querying the Drug–Gene Interaction Database (DGIdb v5.0).^117^

#### Motif deviation, circadian TFs, and TF enrichment network

The single-nucleus TF motif deviation scores were calculated using ArchR’s implementation of chromVAR,^100^ and motif annotations were assigned based on the CIS-BP database. To identify circadian TFs, the single-nucleus TF motif deviation scores were aggregated into a pseudo-bulk matrix across replicates and experimental time points (CT4, CT10, CT16, and CT22) for each cell type. The resulting pseudo-bulk matrices for each type were then analyzed using the DiscoRhythm R package to detect TFs exhibiting 24-hour rhythmic patterns.^31^ The DiscoRhythm workflow applied for circadian TF identification followed the same procedures described for circadian gene detection.

Expression of circadian TFs was then examined in each cell type. TF symbols were converted from protein names to gene symbols using SynGo (https://www.syngoportal.org/convert), and TFs without matched gene symbols were excluded from RNA expression assessment and downstream analyses. Additionally, TFs with no or low expression, defined as those within the lowest 10% of expression among all TFs for each cell type, were removed from further analysis.

Circadian TFs passing these filtering criteria were used to construct TF enrichment networks and listed in Table S3. The single-nucleus TF deviation matrix was subset to include only circadian TFs and transposed to obtain a TF-by-cell matrix. Dimensionality reduction was performed on this matrix using PCA, followed by the calculation of TF neighborhood relationships with the *sc.pp.neighbors* function in Scanpy. Pairwise connectivity and distance metrics among TFs were then exported to Cytoscape for network visualization.^98^ In Cytoscape, each TF node was annotated with a weighted enrichment deviation score specific to each cell type.^98^ This score was calculated by averaging TF deviation scores into a pseudo-bulk matrix across 14 cell types and weighting the values by ln(1/*p*-value), derived from the circadian TF analysis (i.e. pseudo-bulk TF deviation score × ln(1/*p*-value)). Because only TFs with *p*-values less than 0.02 were considered circadian TF in a given type, TFs lacking a *p*-value in a particular type (i.e. not identified as circadian) were assigned a weight of zero (ln(1/*p*-value) = 0).

Circadian TF networks were further constructed specifically for cancer epithelial cells. Only TFs passing the aforementioned filtering steps were included. For this analysis, the TF-by-cell matrix was limited to cancer epithelial cells and their circadian TFs. PCA and neighborhood graph calculations were first performed to capture the relationships among TFs. A partition-based graph abstraction (PAGA) was then generated using the *sc.tl.paga* function in Scanpy,^83^ with the “group” parameter defined by discrete acrophase information. Circadian TFs were categorized into eight acrophase bins (P1–P8) based on their acrophase values. The resulting PAGA graph was subsequently used to initialize the UMAP embedding with the parameter “init_pos=paga”. Visualization of both the UMAP and PAGA graphs was performed using the *sc.pl.paga_compare* function and selected circadian TFs of interest were labeled.

#### Functional enrichment of circadian TF-regulated genes

The TF-regulated cCRE–gene pairs were identified by intersecting the binding sites of selected circadian transcription factors (e.g., *Arntl*, *Nr1d1*, and *Dbp*) with cCRE–gene pairs specific to the cell types enriched for each TF. The resulting TF-regulated circadian genes were subjected to GO enrichment analysis.^97^

#### Copy number variation analysis

Copy number variation (CNV) analysis was conducted using the inferCNV R package (v1.12.0) (https://github.com/broadinstitute/inferCNV) to identify large-scale chromosomal alterations from single-nucleus RNA sequencing data. We used CD45⁻ cell populations, including cancer-associated fibroblasts (CAFs), normal epithelial cells, endothelial cells, perivascular-like (PVL) cells, and cancer epithelial cells. The cells except for cancer epithelial cells were set as reference cells. The inferCNV pipeline was executed with the parameters “cutoff = 0.1, denoise=T, cluster_by_groups=T”, while all other parameters were kept at default settings. To quantify CNV levels for individual cells, the gain and loss expression matrix generated by inferCNV was rescaled to a range between -1 and 1. A CNV score was then computed for each cell as the sum of squared normalized expression deviations across all genes. The resulting CNV scores were visualized by the *Plot_Density_Custom* function from the scCustomize R package (v.2.1.1).^91^

#### CYCLOPS2.0-based circadian phase estimation of human TNBC samples

To estimate the circadian phase of human TNBC samples,^23^ we utilized CYCLOPS 2.0.^55,78^ First, to capture immune-relevant circadian structure while accounting for cell-type specificity, we curated an immune-informed circadian seed gene set. This set was constructed by taking the union of strongly correlated circadian genes identified in mouse CD4^+^ T, CD8^+^ T, and myeloid compartments and mapping them to their human orthologs and listed as Table S5. The normalized pseudo-bulk expression matrix, specifically derived from CD4^+^ T cells, CD8^+^ T cells, and Myeloid cells, served as the input. To account for transcriptional variations driven by cellular identity, cell type labels were incorporated into the CYCLOPS model as covariates. The algorithm was then executed to infer circadian phase angles for each cell type within each sample. To define a robust patient-level circadian phase, referred to as the Donor Internal Phase (DIP), the inferred phases from distinct immune compartments were aggregated using a circular mean. To ensure robustness against noise, an iterative outlier-removal strategy was applied, excluding cell types whose phases deviated by more than π/2 from the current mean. The mean was then iteratively recalculated until no further outliers were detected or a maximum of five iterations was reached. The final DIPs were subsequently used to assign samples to discrete circadian time groups for downstream analyses. To align circadian phases between nocturnal mice and diurnal humans, a 12-hour phase shift was applied to the predicted human DIPs.

For circadian rhythmicity analysis, samples were subsequently assigned to six circadian phase groups based on their inferred DIPs. The normalized expression matrix was aggregated into a pseudo-bulk matrix across biological replicates and predicted human DIPs for each cell type, followed by rhythmic gene detection using the DiscoRhythm R package as described above.^31^

