## Supplementary Figures for "Circadian misalignment underlies immune escape in breast cancer"

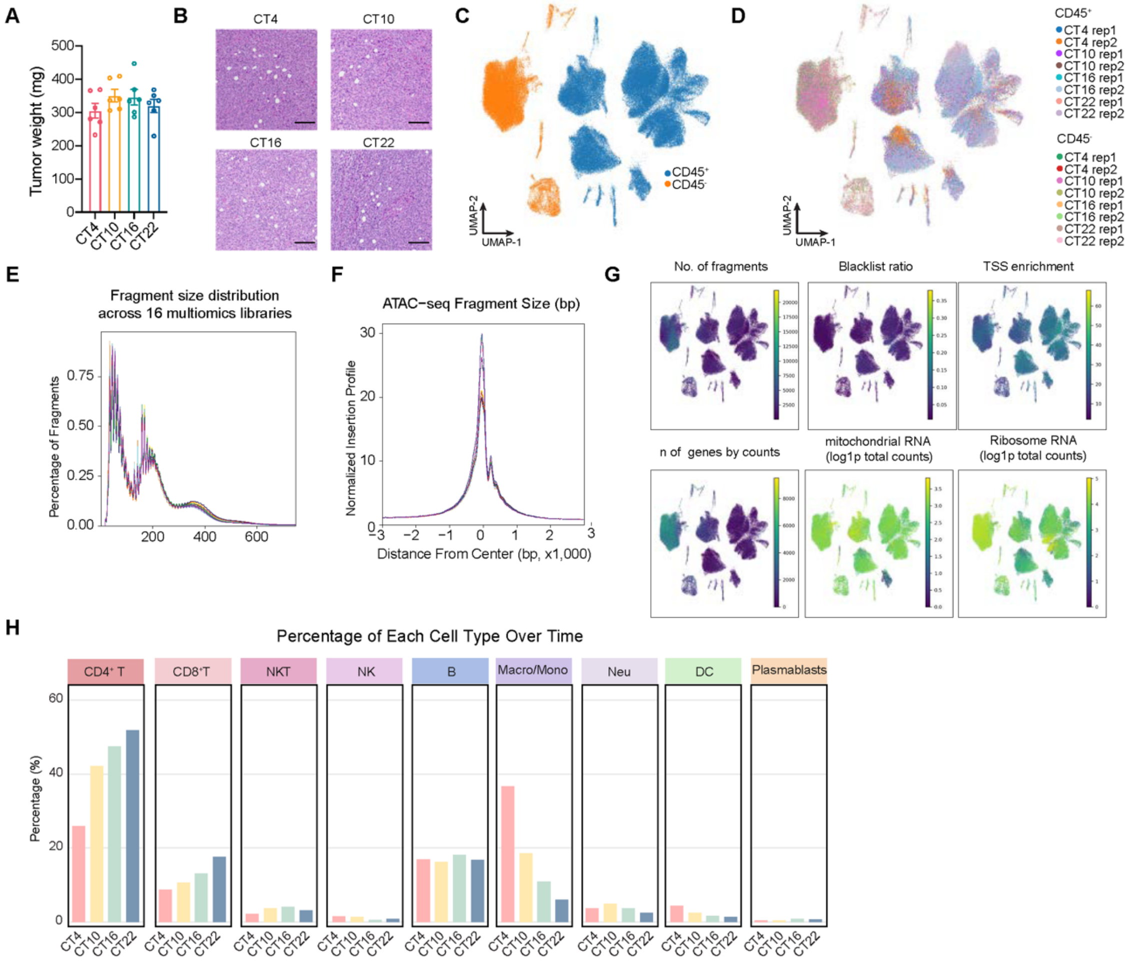


**Figure S1. Experimental design, data quality assessment, and cell-type composition of tumor single-nucleus multiomic profiles, related to Figure 1.**

(A) Bar plots showing the tumor weights collected from four different timepoints. *n*=6 mice per group. No significant difference among groups (one-way ANOVA, ns).

(B) Representative hematoxylin and eosin (H&E) staining of tumor tissues collected at four circadian time points.

(C) UMAP visualizations of snMultiome-seq profiles, with nuclei colored by CD45 status (CD45$^+$ versus CD45$^-$).

(D) UMAP visualizations of snMultiome-seq profiles, with cells colored by 16 libraries.

(E) Fragment size distribution showing high data quality across all 16 snMultiome libraries.

(F) Transcription start site (TSS) enrichment profiles of snMultiome-seq data across 16 snMultiome libraries, indicating strong signal-to-noise ratios.

(G) Quality-control metrics of snMultiome-seq projected onto the UMAP embedding, including the number of ATAC fragments, blacklist ratio, TSS enrichment, number of detected genes, mitochondrial RNA count (log1p total counts), and ribosomal RNA count (log1p total counts) per nucleus.

(H) Bar plots showing the percentage of each immune cell type (CD4$^+$ T, CD8$^+$ T, NKT, NK, B, Macro/Mono, Neu, DC, Plasmablasts) across four time points.

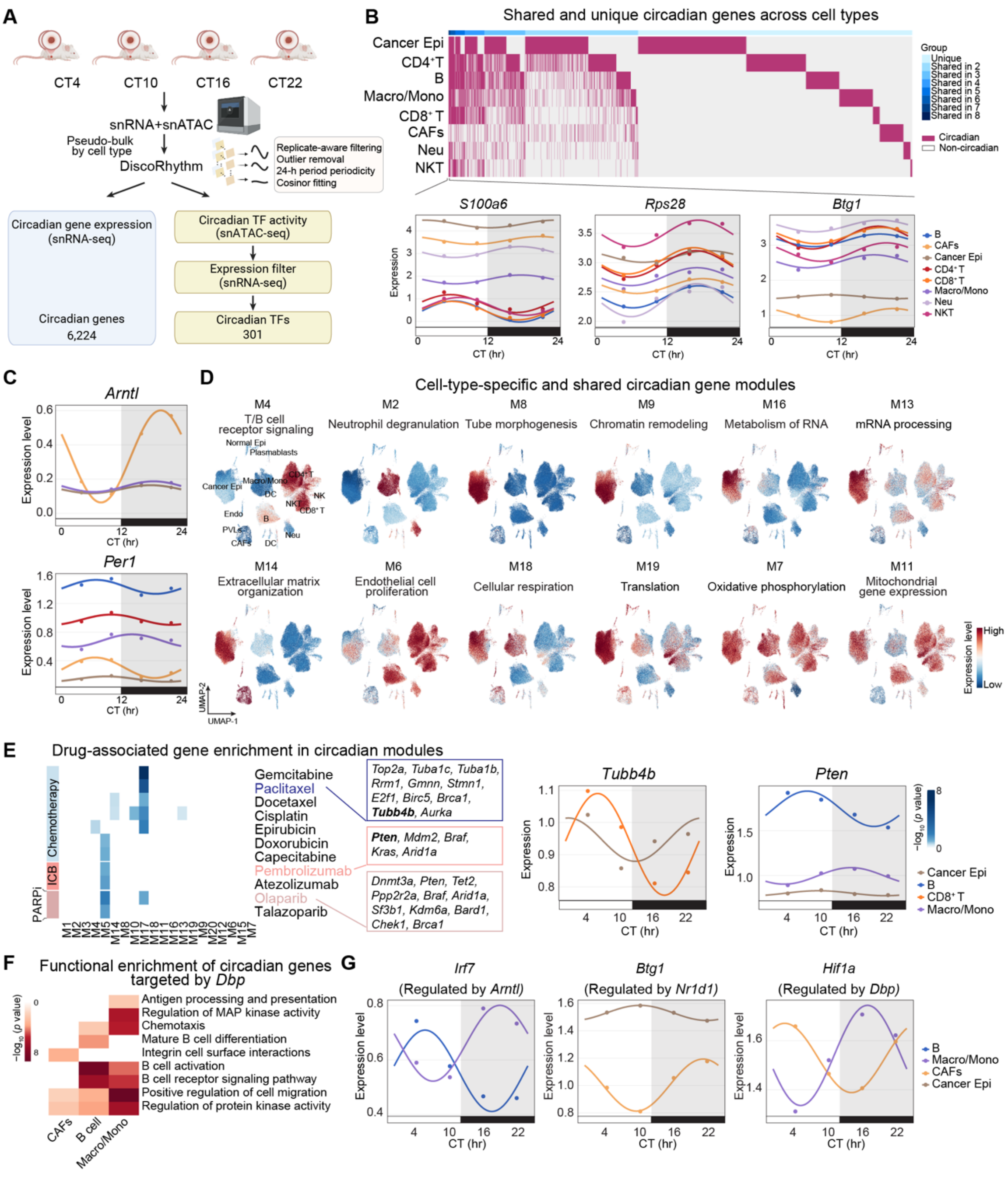


**Figure S2. Circadian transcriptomic and transcription factor networks across TME cell types, related to Figure 2.**

(A) Schematic overview of the analysis pipeline. Circadian gene expression (snRNA-seq) and transcription factor (TF) motif activity (snATAC-seq) were identified across four circadian time points (CT4, CT10, CT16, and CT22) using DiscoRhythm.

(B) Top: Heatmap showing shared and unique circadian genes among major cell types. Each column represents a rhythmic gene, with colors indicating whether it is unique to one cell type or shared among multiple cell types (light to dark blue, representing genes shared in 2–8 cell types). Circadian genes are shown in purple, and non-circadian genes in light gray. Bottom: Representative oscillatory expression patterns of circadian genes shared across eight cell types (*S100a6*, *Rps28*, and *Btg1*). Shaded areas indicate the dark phase (CT12–CT24/CT0).

(C) Representative oscillatory expression patterns of core circadian clock genes (*Arntl* and *Per1*) across indicated cell types. Shaded areas indicate the dark phase (CT12–CT24/CT0).

(D) UMAP visualizations of selected circadian gene modules. The color bar indicates the module expression level, ranging from high (deep red) to low (deep blue).

(E) Left: Heatmap showing the enrichment of drug associated genes within circadian gene modules. Several therapeutic agents, including chemotherapy, immune checkpoint blockade (ICB) and PARP inhibitors (PARPi), are linked to rhythmic pathways. The color bar indicates the $-\log_{10}(p$ value) of enrichment, ranging from high (deep blue) to low (white). Right: Rhythmic expression of representative drug-associated genes (*Tubb4b* and *Pten*) across multiple cell types. Shaded areas indicate the dark phase (CT12–CT24/CT0).

(F) Heatmaps showing the functional enrichment of circadian genes regulated by core circadian clock TF *Dbp* across selected cell types. The color bar indicates the $-\log_{10}(p$ value) of enrichment, ranging from high (deep red) to low (white).

(G) Rhythmic expression of selected core clock component-regulated genes (*Irf7, Btg1* and *Hif1a*) across multiple TME cell types, exhibiting cell-type–specific phase differences. Shaded areas indicate the dark phase (CT12–CT24/CT0).

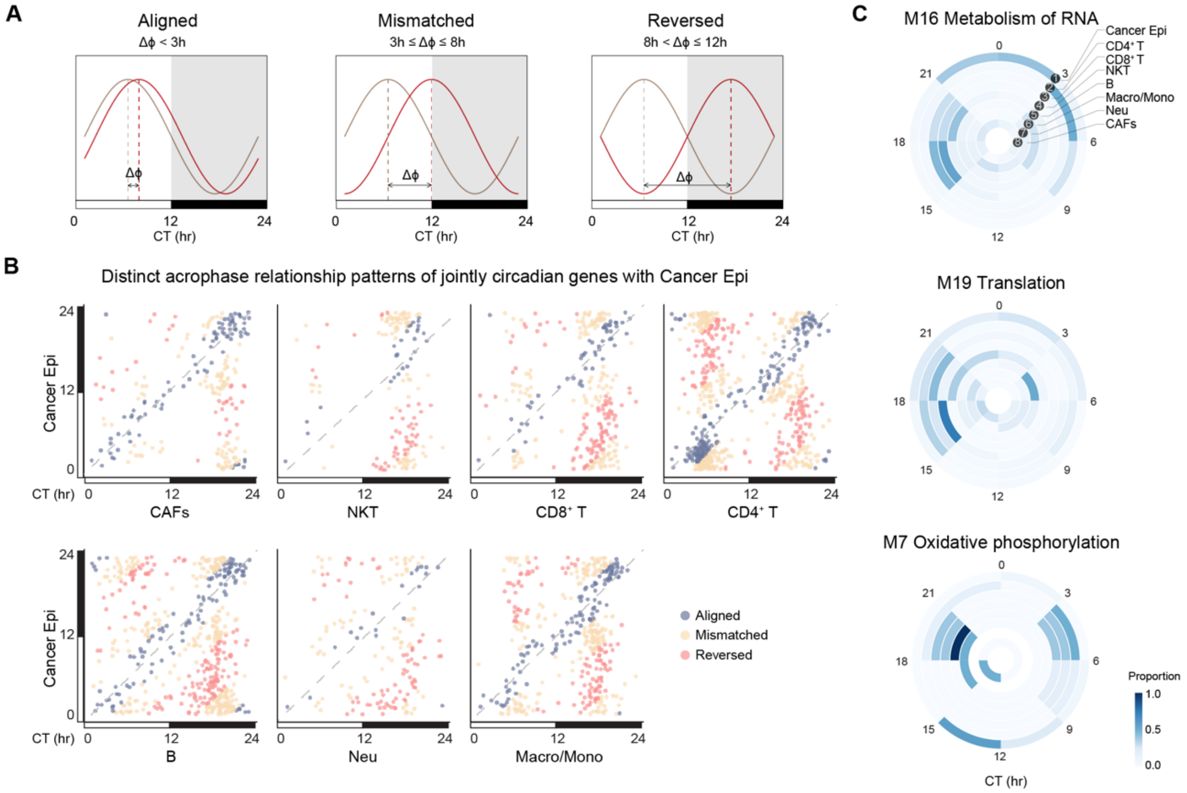


**Figure S3. Circadian phase comparison and module-specific phase distribution across cell types, related to Figure 3.**

(A) Schematic illustration of three circadian phase relationship categories across cell types defined by acrophase difference (Δϕ): aligned (< 3 h), mismatched (3–8 h), and reversed (8–12 h).

(B) Scatter plots showing the acrophase of joint circadian genes between cancer epithelial cells and other major cell types. Each dot represents one shared circadian gene. Genes are classified as aligned (Δϕ <3 h, blue), mismatched (Δϕ 3-8 h, yellow), or reversed (Δϕ 8–12 h, red). The dashed line indicates perfect phase alignment.

(C) Circular plots showing the acrophase distribution of genes within M16 (top), M19 (middle) and M7 (bottom) circadian modules across different cell types. Color intensity indicates the proportion of circadian genes peaking within each of the eight phase bins (P1–P8), from high (dark blue) to low (white).

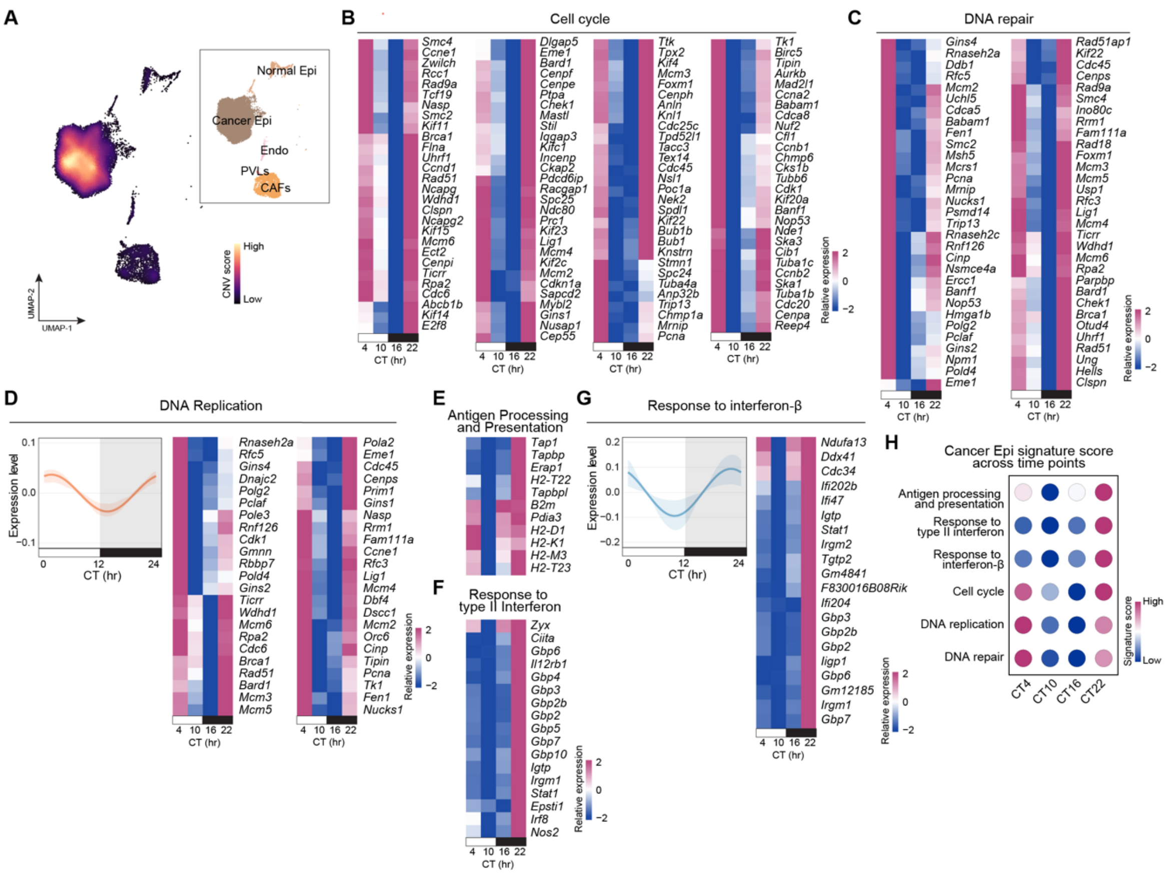


**Figure S4. Circadian regulation in cancer epithelial cells, related to Figure 4.**

(A) UMAP visualization of inferred copy-number variation (CNV) scores in $CD45^{-}$ cells (left) and corresponding cell-type annotations (right). The color bar indicates CNV scores from high (bright yellow) to low (deep purple).

(B-G) Heatmaps showing rhythmic expression patterns of circadian genes involved in cell cycle (B), DNA repair (C), DNA replication (D), antigen processing and presentation (E), response to type II interferon (F), and response to interferon-β (G) in cancer epithelial cells across circadian time points. The corresponding line plots for panels D and G are shown to the left of the heatmaps, whereas those for panels B, C, E, and F are shown in Figure 4B. Bold lines represent the mean fitted circadian curve across all genes involved in each pathway, and shaded areas indicate the interquartile range (25th–75th percentile). Gray areas indicate the dark phase (CT12–CT24/CT0).

(H) Bubble plot showing pathway signature score dynamics across circadian time points (CT4–CT22). The color bar indicates the average signature score, ranging from high (magenta) to low (blue).

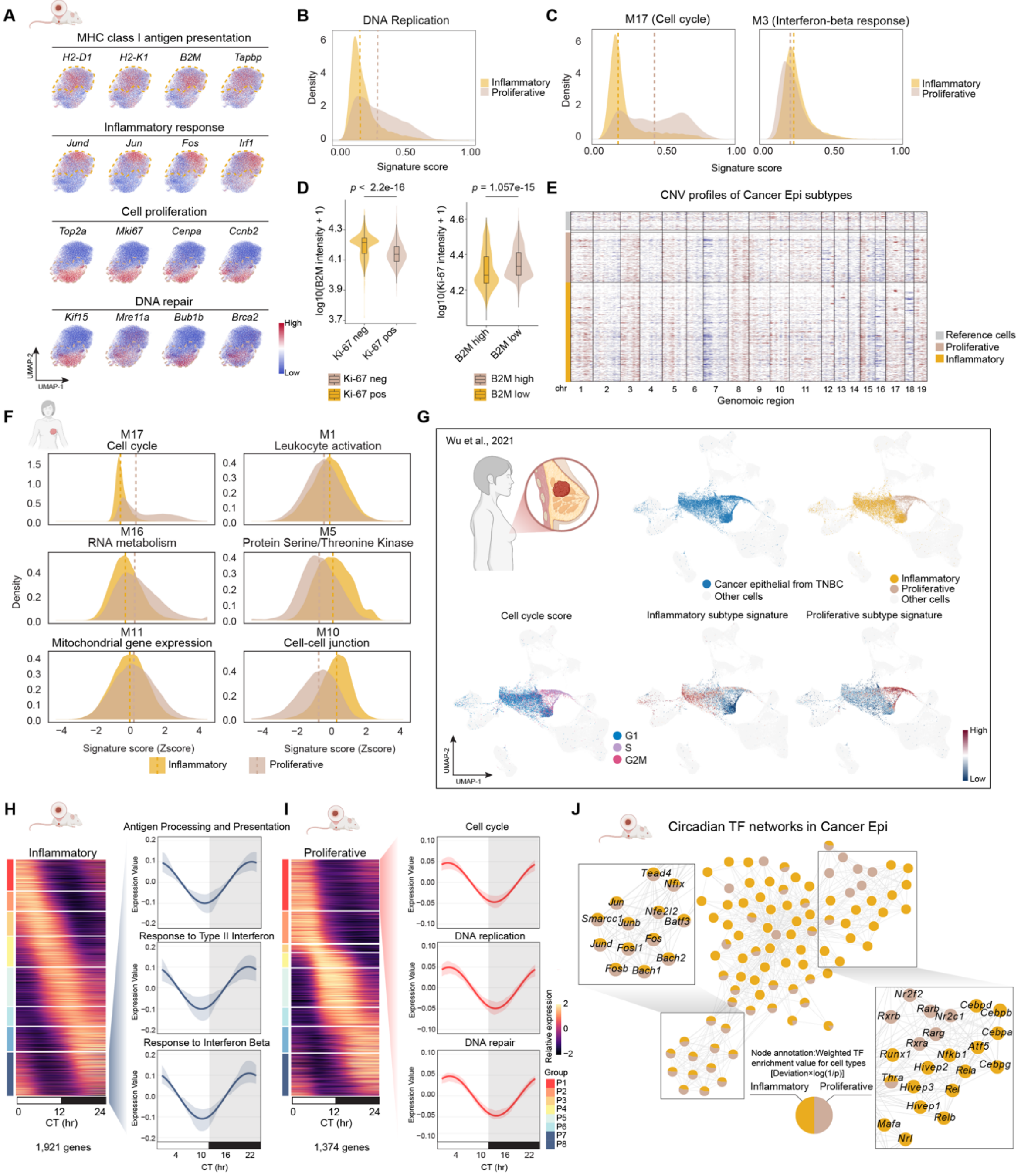


**Figure S5. State-specific circadian transcriptional programs and transcription factor networks in cancer epithelial cells, related to Figure 4.**

(A) UMAPs showing the expression of representative genes associated with MHC class I antigen presentation, inflammatory response, cell proliferation, and DNA repair in cancer epithelial cells. The color bar indicates the expression level from high (red) to low (blue).

(B) Density plots showing the distribution of DNA replication signature scores across inflammatory and proliferative cancer epithelial cells in mouse tumors.

(C) Density plots showing the distribution of circadian module scores (M17, cell cycle; M3, interferon-β response) across inflammatory and proliferative cancer epithelial cells in mouse tumors.

(D) Violin plots showing reciprocal expression of B2M and Ki-67 in cancer epithelial cells. Left, B2M intensity in Ki-67 negative versus Ki-67 positive cancer epithelial cells. Right, Ki-67 intensity in B2M high versus B2M low cancer epithelial cells. Boxes indicate median and interquartile range; *p* values were calculated using a two-sided Wilcoxon rank-sum test.

(E) Single-cell inferred CNV profiles of inflammatory and proliferative cancer epithelial cells across genomic regions, with non-cancer epithelial cells shown for reference.

(F) Density plots showing selected circadian module signature distributions across inflammatory and proliferative cancer epithelial cells based on reanalysis of published human TNBC single-cell data.[49]

(G) UMAP visualizations of reanalyzed human TNBC single-cell RNA-seq data,[28] highlighting cancer epithelial cells (top left) and their subtype classification into inflammatory and proliferative states (top right). Bottom panels show cell-cycle phase scores and subtype signature scores from mouse data (inflammatory and proliferative) across epithelial populations. Color scale represents the expression level from high (red) to low (blue).

(H-I) Heatmap showing rhythmic expression patterns of circadian genes in inflammatory (H) and proliferative (I) cancer epithelial cells. Representative pathways enriched in each subtype (right) exhibit distinct temporal expression patterns, with immune-associated programs (antigen processing and presentation, interferon responses) in inflammatory cancer epithelial cells mainly peaking at phase P8, and proliferative programs (cell cycle, DNA replication, DNA repair) in proliferative cancer epithelial cells predominantly peaking at phase P1. Lines represent the mean circadian fitting curve across pathway genes; shaded areas indicate the interquartile range (25th–75th percentile), and gray area indicate the dark phase (CT12–CT24/CT0).

(J) Circadian TF networks in cancer epithelial cells inferred from motif binding with circadian rhythmicity. Each node represents a TF, colored by inflammatory or proliferative cancer epithelial cells, and scaled by TF deviation score weighted by circadian rhythmicity.

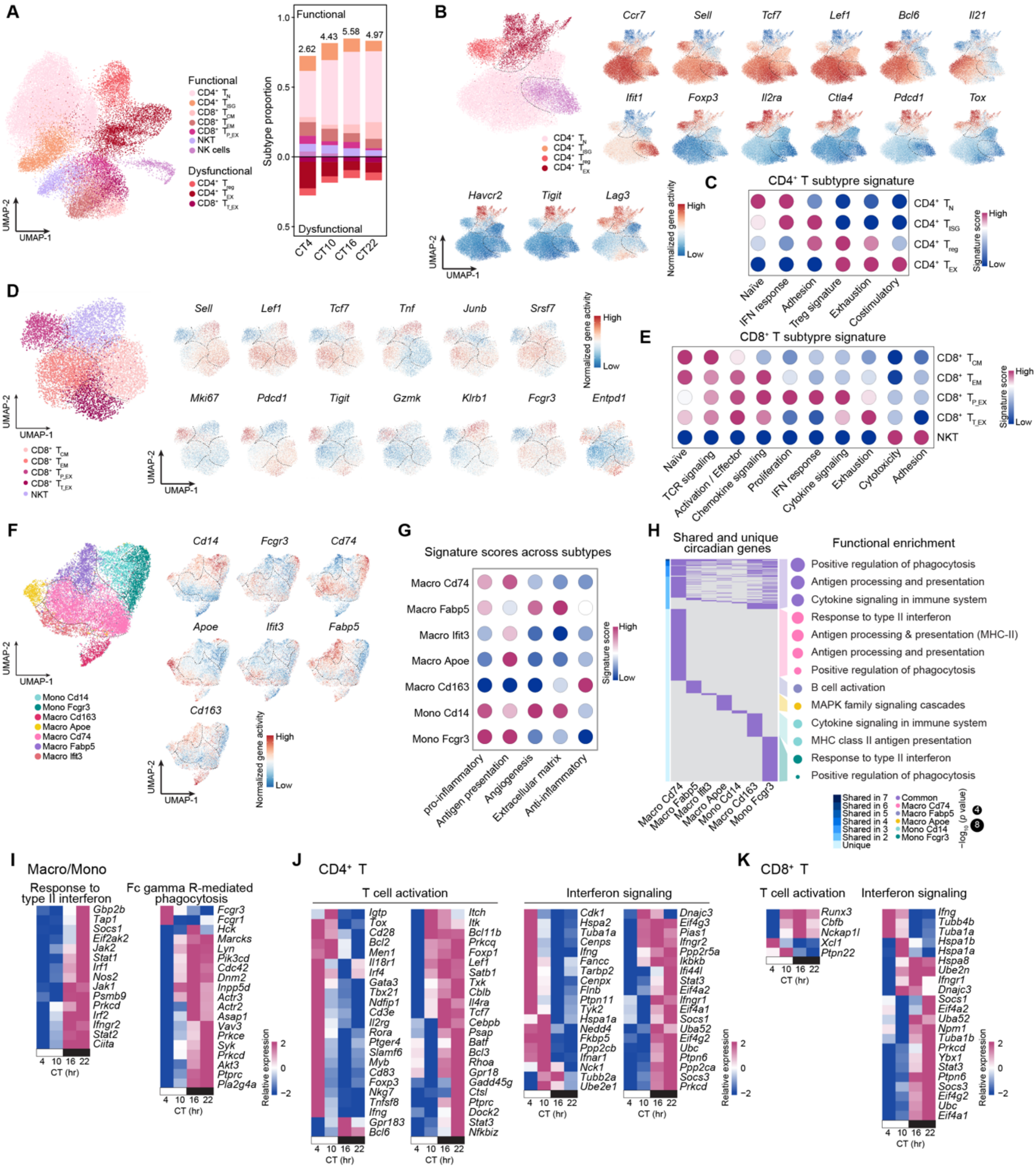


**Figure S6. Cell-type-specific circadian regulation of immune activation in CD4$^+$ T, CD8$^+$ T and Macro/Mono subtypes, related to Figure 5.**

(A) UMAP visualization showing major T and NK cell subtypes, including CD4$^+$ and CD8$^+$ T cell subsets, NKT, and NK cells (left). Diverging stacked bar plot (right) showing the relative proportions of functional states (NK, NKT, CD4$^+$T$_{N}$, CD4$^+$T$_{ISG}$, CD8$^+$T$_{CM}$, CD8$^+$T$_{P_EX}$) and dysfunctional states (CD4$^+$T$_{reg}$, CD4$^+$T$_{EX}$, CD8$^+$T$_{T_EX}$) across circadian time points. Values above bars indicate the functional-to-dysfunctional ratio at each time point.

(B) UMAPs displaying representative marker gene activity levels inferred from snATAC-seq data for CD4$^+$ T cell subsets. The color bar indicates gene activity level from high (red) to low (blue).

(C) Bubble plot showing CD4$^+$ T cell subtype signature scores. The color bar indicates the average signature score, ranging from high (magenta) to low (blue).

(D) UMAP visualization showing CD8$^+$ T cell subsets (left). UMAPs on the right display representative marker gene activity levels inferred from snATAC-seq data for CD8$^+$ T cell subsets. The color bar indicates gene activity level from high (red) to low (blue).

(E) Bubble plot showing CD8$^+$ T cell subtype signature scores. The color bar indicates the average signature score, ranging from high (magenta) to low (blue).

(F) UMAP visualization showing Macro/Mono subtypes. representative marker gene activity levels inferred from snATAC-seq data for Macro/Mono subsets. The color bar indicates gene activity level from high (red) to low (blue).

(G) Bubble plot showing Macro/Mono subtypes signature scores. The color bar indicates the average signature score, ranging from high (magenta) to low (blue).

(H) Heatmap showing shared and unique circadian genes across macrophage and monocyte (Macro/Mono) subtypes, with circadian genes indicated in purple. Color intensity from light to dark blue represents the degree of gene sharing, ranging from unique to shared across all seven subtypes. The corresponding GO enrichment (right) highlights representative biological processes associated with each group. Dot size indicates $-\log_{10}(p \text{ value})$ of pathway enrichment, and dot color represents the corresponding subtype category (Common, Macro Cd74, Macro Fabp5, Macro Apoe, Mono Cd14, or Mono Fcgr3).

(I–K) Heatmaps showing rhythmic expression patterns of circadian genes involved in key immune pathways, including type II interferon response and Fc gamma R-mediated phagocytosis in Macro/Mono (I), and T cell activation and interferon signaling in $CD4^+$ T cells (J) and $CD8^+$ T cells (K). The color bar represents expression levels ranging from high (magenta) to low (blue). Related to Figure 5G.

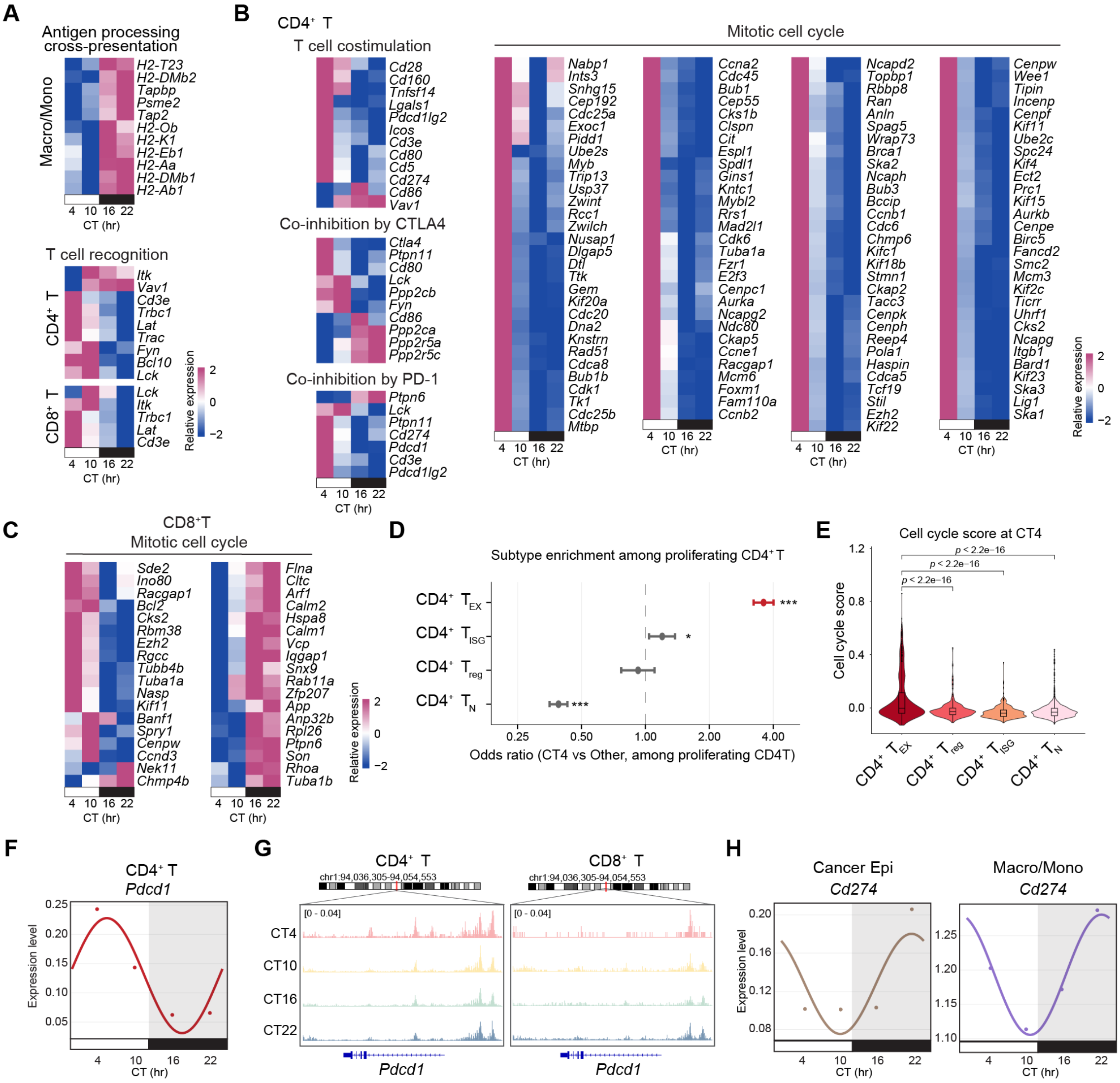


**Figure S7. Circadian regulation of T cell recognition, proliferation, and immune checkpoint inhibition dynamics, related to Figure 6.**

(A) Heatmaps showing rhythmic expression patterns of circadian genes involved in key immune pathways, including antigen processing and cross-presentation in Macro/Mono and T cell recognition in CD4$^+$ T and CD8$^+$ T. The color bar represents expression levels ranging from high (magenta) to low (blue). Related to Figure 6A.

(B) Heatmaps showing rhythmic expression patterns of circadian genes involved in key immune pathways in CD4$^+$ T, including T cell costimulation, co-inhibition by CTLA4/PD-1, and mitotic cell cycle. The color bar represents expression levels ranging from high (magenta) to low (blue). Related to Figure 6D.

(C) Heatmaps showing rhythmic expression patterns of circadian genes involved in mitotic cell cycle in CD8$^+$ T. The color bar represents expression levels ranging from high (magenta) to low (blue). Related to Figure 6D.

(D) Forest plot showing subtype enrichment among proliferating CD4$^+$ T cells at CT4 compared with other circadian time points. Odds ratios indicate preferential enrichment of specific CD4$^+$ T subtypes during the proliferative phase. Statistical significance is denoted as * $p < 0.05$, ** $p < 0.01$, *** $p < 0.001$.

(E) Violin plots showing cell cycle scores of CD4$^+$ T subtypes at CT4. Box boundaries represent interquartile ranges, center lines denote medians, and whiskers indicate data range. Statistical significance was assessed using the Kruskal–Wallis test.

(F) Line plots showing circadian fitting curves of *Pdcd1* in CD4$^+$ T cells. Dots represent mean expression values at each circadian time point. The gray area indicates the dark phase (CT12–CT24/CT0).

(G) Chromatin accessibility profiles showing time-of-day–dependent changes of the *Pdcd*1 locus across a day in $CD4^+$ and $CD8^+$ T cells.

(H) Line plots showing circadian fitting curves of *Cd274* (PD-L1) in Cancer Epi (left) and Macro/Mono (right). Dots represent mean expression values at each circadian time point. The gray area indicates the dark phase (CT12–CT24/CT0).

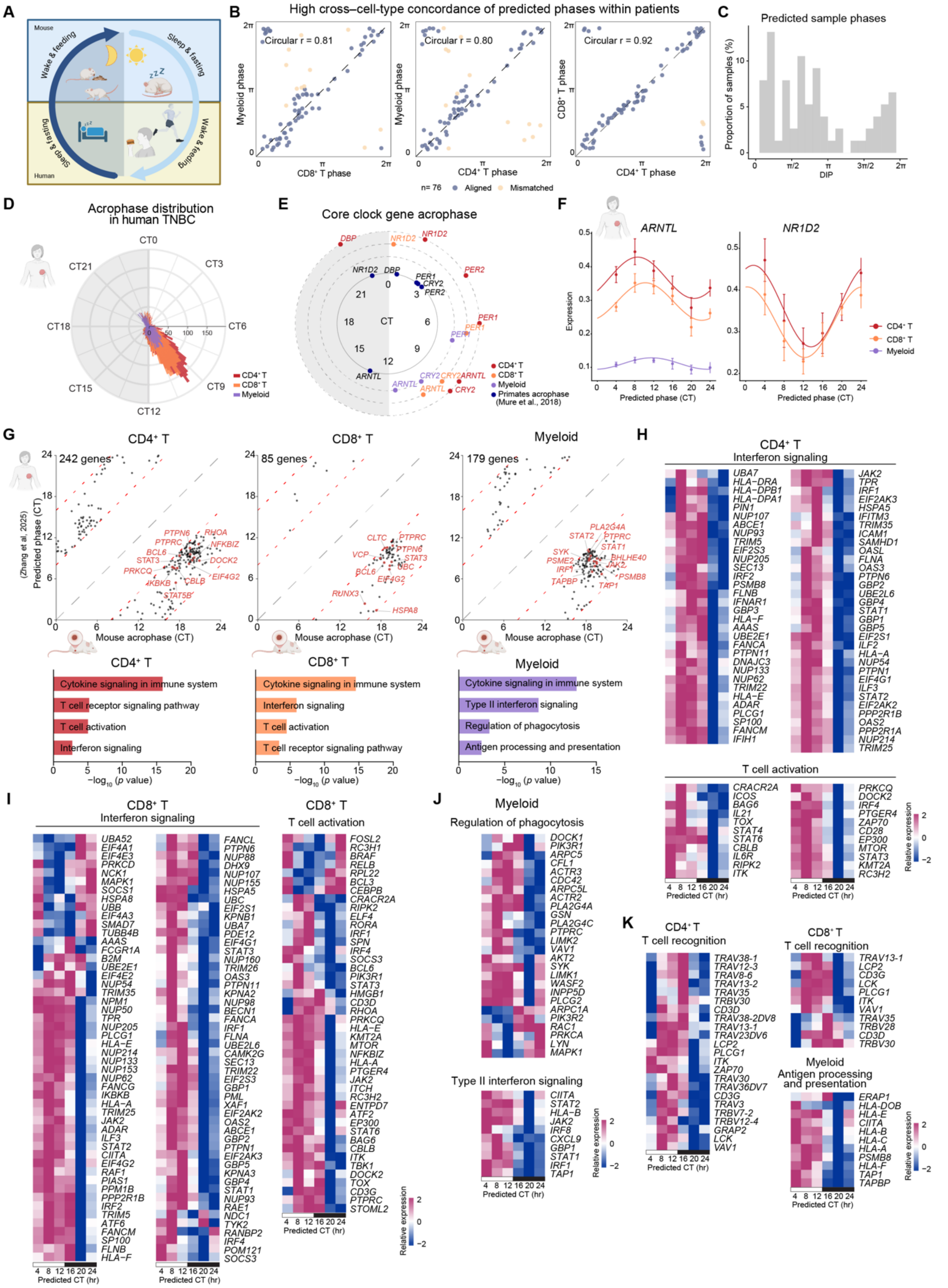


**Figure S8. Circadian regulation of CD4⁺ T, CD8⁺ T and myeloid cells in human TNBC, related to Figure 7.**

(A) Schematic illustrating the inverted circadian organization between nocturnal mouse models and diurnal humans, highlighting the expected phase reversal between species.

(B) Pairwise comparisons of inferred circadian phases (φ) between immune cell types within the same patient, showing high cross-cell-type concordance. Scatter plots display myeloid versus CD8⁺ T cell phases (left), myeloid versus $CD4^+$ T cell phases (middle), and $CD8^+$ versus $CD4^+$ T cell phases (right).

(C) Distribution of inferred patient-level circadian phases (DIP) across all human TNBC samples, showing a non-uniform distribution with enrichment in specific circadian intervals.

(D) Polar plot showing the distribution of inferred acrophases of circadian genes across $CD4^+$ T cells, $CD8^+$ T cells and myeloid in human TNBC.

(E) Circular comparison of acrophase of core clock genes across immune cell types in human TNBC, overlaid with reported acrophase from diurnal primates. Relative phase ordering is preserved, supporting the validity of human circadian phase transformation.

(F) Circadian fitting curves for representative core clock genes (*ARNTL* and *NR1D2*) in $CD4^+$ T cells, $CD8^+$ T cells, and myeloid cells. Points represent mean expression at each inferred circadian time; error bars indicate SEM.

(G) Top: Scatter plots comparing acrophases of circadian genes showing conserved but phase-reversed regulation between mouse tumors and human TNBC in $CD4^+$ T cells (left), $CD8^+$ T cells (middle), and myeloid cells (right). Each point represents one gene; immune-related genes are highlighted. Numbers indicate the total number of such genes in each cell type. Bottom: Functional enrichment of these genes, highlighting immune activation pathways in T cells and phagocytosis and antigen presentation in myeloid cells.

(H and I) Heatmaps showing rhythmic expression patterns of circadian genes involved in interferon signaling and T cell activation in $CD4^+$ T cells (H) and $CD8^+$ T cells (I), related to Figure 7D and 7G. The color bar represents expression levels ranging from high (magenta) to low (blue).

(J) Heatmaps showing rhythmic expression patterns of circadian genes involved in regulation of phagocytosis and type II interferon signaling in myeloid cells, related to Figure 7J. The color bar represents expression levels ranging from high (magenta) to low (blue).

(K) Heatmaps showing rhythmic expression patterns of circadian genes involved in T cell recognition in $CD4^+$ T and $CD8^+$ T cells and antigen processing and presentation in myeloid, related to Figure 7L. The color bar represents expression levels ranging from high (magenta) to low (blue).
